# Adaptive plasticity of aspartate metabolism in succinate dehydrogenase-deficient cancer cells

**DOI:** 10.64898/2026.05.18.726122

**Authors:** David Sokolov, Eric Zheng, Serwah Danquah, Madeleine L. Hart, Lucas B. Sullivan

## Abstract

Succinate dehydrogenase (SDH) supports cancer cell proliferation by enabling oxidative biosynthesis of the amino acid aspartate, yet SDH loss can also drive tumorigenesis. To cope with SDH loss, cancer cells can engage alternative aspartate synthesis pathways; however, the variables dictating pathway usage and adaptive mechanisms involved are incompletely understood. Here, we systematically profile the adaptation of SDH-knockout cancer cells and find that cells can adapt to SDH loss via at least two distinct mechanisms: suppression of respiratory complex I or upregulation of pyruvate carboxylase. Each route gives rise to distinct metabolic states with both shared and unique dependencies, but either route allows cells to overcome aspartate limitation, improve proliferative fitness, and mitigate pyrimidine-dependent replication stress. Overall, this work provides a comprehensive view of adaptive aspartate synthesis in SDH-deficient cancer cells, highlights a remarkable redox-constrained metabolic plasticity, and nominates potential metabolic vulnerabilities likely to be shared among SDH-deficient cancer cells.

---

Cancerous cells possess metabolic behaviors distinct from non-cancerous cells ^1^. Some of these metabolic differences reflect emergent consequences of oncogenic programs, including signaling-specific changes to metabolic pathways and flux changes arising from the specific metabolic demands of driving biomass synthesis and cell proliferation ^2,3^. Beyond these cell state-intrinsic effects, further metabolic changes can occur as cancer cells progressively adapt to optimize fitness or mitigate stressors associated with harsh or austere tumor microenvironments ^4^, metastasis ^5–7^, or immune surveillance ^8^. While it’s now widely appreciated that these collective metabolic differences are meaningful for cancer progression ^9^, we know less about which metabolic adaptations are functionally relevant in which cancer contexts, or the nature of the adaptive processes involved.

The development of myriad sequencing-based methods over the past several decades has enabled unprecedented insight into genetic and epigenetic tumor evolution ^10^, but these technologies face challenges in dissecting tumor metabolic adaptation—not only because they only indirectly measure metabolism, but also because the specific selective pressures driving adaptation are often inferred and difficult to verify. Understanding context-specific metabolic adaptations of cancer cells is important, since adapted states may further differentiate tumor from non-tumor tissue and be therapeutically actionable ^11^. Finally, a better understanding of the features of cancer metabolic adaptation may inform responses to other instances of tumor evolution, including acquired treatment resistance ^12^.

One specific context of cancer metabolic adaptation is that where loss of core metabolic enzymes drives tumorigenesis: a prototypical example concerns the enzyme succinate dehydrogenase (SDH), a conserved redox-active enzyme at the heart of mitochondrial and central carbon metabolism. SDH presents a seeming paradox, in that SDH function both supports cancer cell proliferation in part by enabling the synthesis of the amino acid aspartate ^13–15^, but also that SDH impairment promotes several human cancer types via a classical tumor suppressor mechanism ^16^. One defining metabolic feature of SDH deficiency is a dramatic accumulation of the SDH substrate succinate, an ‘oncometabolite’ thought to promote tumorigenesis by interfering with cellular oxygen sensing and gene regulatory pathways, among other things ^17,18^.

Thus, while it is currently thought that SDH loss causes cancer by precipitating a metabolic phenotype conducive to malignant transformation, the mechanisms by which SDH-deficient cancer cells cope with the metabolic consequences of losing SDH are comparatively less understood. Concerning this question, several studies including ours have found that SDH-deficient human and mouse cells can compensate for the loss of oxidative TCA activity by employing alternative metabolic pathways to partially restore aspartate biosynthesis and proliferation ^13–15^. Notably, these studies found evidence of at least two pathways which SDH-deficient cells could use to synthesize aspartate: a glucose-derived pathway requiring pyruvate carboxylase (PC)-mediated anaplerosis and a reductive glutamine-derived pathway previously shown to be active in respiration-impaired cancer cells ^13–15,19,20^.

We recently described a phenomenon whereby SDH-deficient cancer cells can adapt by suppressing the activity of respiratory complex I (CI), which reduces mitochondrial NAD^+^/NADH to drive these two alternative aspartate synthesis pathways ^15^. These results conceptually fit other reports of CI suppression playing a role in SDH-deficient cell metabolic fitness ^13,21^, but stood at odds with studies that characterized distinct SDH-deficient cell systems with apparently intact CI activity ^22,23^. Collectively, these studies reveal a considerable metabolic heterogeneity across systems that may allow SDH-deficient cancer cells to tolerate or even overcome the aspartate limitation caused by SDH loss. However, the variables governing the engagement of specific alternative aspartate synthesis pathways in SDH-deficient cells remain unclear.

Here, we address these questions by measuring the progressive adaptation of a collection of SDHB-knockout clones. In addition to the CI-dependent adaptive route we previously characterized, we find a distinct, CI-independent adaptive trajectory by which SDH-deficient cells can overcome aspartate limitation. Characterization of alternative aspartate synthesis in both CI-suppressed and CI-intact modes reveals distinct metabolic states with both shared and distinct metabolic dependencies. We further identify PC upregulation as one distinct mechanism by which cells can adapt via the CI-independent route, and we find evidence that PC expression influences sensitivity to SDH impairment across cancer cell lines. Finally, we show that adaptation via either trajectory ameliorates pyrimidine deficiency, cell cycling defects, and sensitivity to ATR inhibition ^24^. Overall, our results unify previously disparate findings under a comprehensive model of adaptive aspartate synthesis in SDH-deficient cells, which advances efforts to characterize and target the unique metabolism of SDH-deficient tumors, as well as other instances in which canonical aspartate production is constrained.

## Results

### Complex I-independent adaptation of SDH-deficient cancer cells

To rigorously evaluate the kinetics and reproducibility of adaptation in response to SDH deficiency, we used CRISPR/Cas9 to knock out the essential SDH subunit SDHB in 143B osteosarcoma cells and isolated five clonal cell populations harboring no SDHB expression (Figure 1A-B). Consistent with previous findings ^13–15^, these SDH-deficient clones require exogenous pyruvate to synthesize aspartate and proliferate (Figure S1B). However, even in pyruvate-replete culture media, these cells experience aspartate limitation, since aspartate supplementation further improves their proliferation rates by 1.5-2 fold (Figure 1C, S1B). To verify that our knockout strategy was on-target, we established SDHB ‘addback’ clones by stably re-expressing V5-tagged SDHB in three of these clones (Figure S1A). Addback clones proliferated at rates typical for 143B cells ^15^ regardless of pyruvate or aspartate supplementation (Figure S1B), verifying that the pyruvate auxotrophy and aspartate limitation of SDHB-KO clones are metabolic phenotypes stemming from SDH deficiency.

**Fig. 1.**
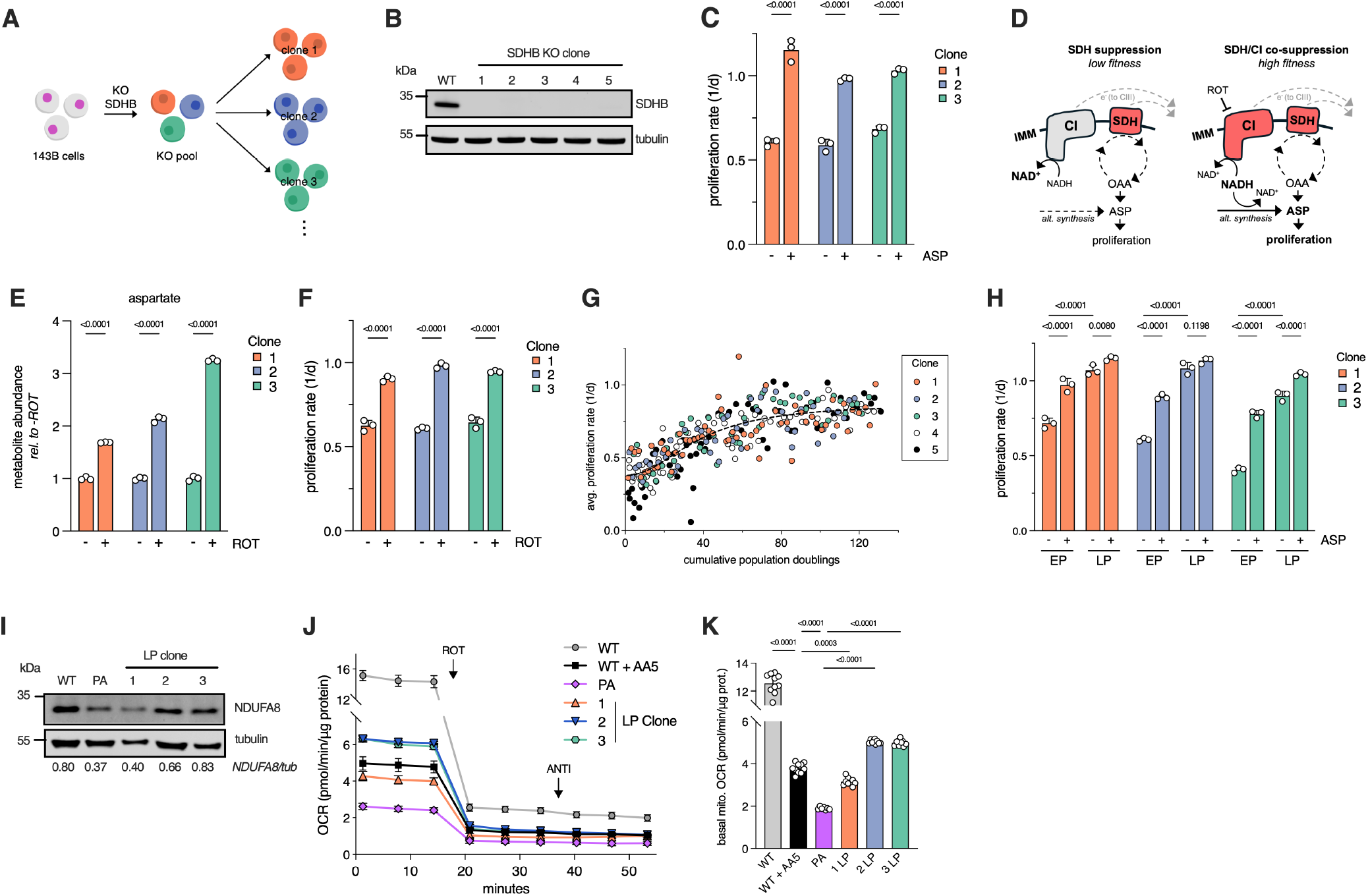
Complex I-independent adaptation of SDH-deficient cancer cells. (**A**) Schematic illustrating the generation of clonal SDHB-knockout 143B cells (**B**) Representative western blot showing levels of SDHB and tubulin loading control in wild-type parental cells (WT) and SDHB knockout clones 1-5 (**C**) Absolute proliferation rates (mean +/-S.D.) of SDHB KO clones 1-3 with or without 20 mM aspartate supplementation. (n=3) (**D**) Schematic illustrating the effects of CI/SDH co-suppression ^15^ on alternative aspartate synthesis and cell fitness. (**E**) Relative whole-cell aspartate levels (mean +/-S.D.), measured using LCMS, in SDHB-KO clones 1-3 after 6 hours of treatment with vehicle control or 50 nM rotenone (ROT). Levels are normalized to the vehicle-treated condition in each respective clone. (n=3) (**F**) Absolute proliferation rates (mean +/-S.D.) of SDHB KO clones 1-3 with or without 50nM rotenone (ROT) supplementation. (n=3) (**G**) Average inter-passage proliferation rates (see methods) of five SDHB-knockout clones over 120 cumulative population doublings. See Figure S1 for plots highlighting individual clones. (**H**) Absolute proliferation rates (mean +/-S.D.) of early passage (EP) and late passage (LP) clones 1-3 with or without 20 mM aspartate supplementation. (n=3) (**I**) Representative western blot showing levels of NDUFA8 and tubulin loading control in wild-type parental cells (WT), a previously-adapted (PA) SDHB-KO clone that adapted by suppressing complex I ^15^, and late passage (LP) SDHB knockout clones 1-3. Normalized NDUFA8 band densities are shown below each respective lane. (**J**) Normalized oxygen consumption rate (OCR) traces (mean +/-S.D.) for wild-type parental cells (WT), parental cells treated with 5 *µ*M Atpenin A5 (WT + AA5), PA, and early passage (EP) and late passage (LP) SDHB knockout clones 1-3. Injections of rotenone (ROT) and antimycin (ANTI) are shown with arrows. (n=7-12) (**K**) Basal mitochondrial oxygen consumption rates (OCR) (mean +/-S.D.) quantified from data in (**J**). (n=7-12). Statistical significance determined using an ordinary two-way ANOVA and uncorrected Fisher’s LSD with single pooled variance (**C, E-F**) or ordinary one-way (**K**) or two-way (**H**) ANOVA and Tukey’s multiple comparisons test with a single pooled variance.

We previously reported that SDH-deficient cancer cells can adapt by downregulating respiratory complex I (CI), which reduces mitochondrial NAD^+^/NADH to drive alternative aspartate synthesis necessary for maximal proliferation ^15^ (Figure 1D). To verify that CI suppression similarly benefits these SDHB-KO cells, we treated KO clones 1-3 with the CI inhibitor rotenone and measured the resulting metabolic and functional effects using liquid chromatography-mass spectrometry (LCMS) and proliferation assays. Consistent with our previous results, rotenone reduced whole-cell NAD^+^/NADH, increase whole-cell aspartate levels, and improved proliferation to a similar degree as aspartate supplementation (Figure S1C, 1E-F).

These results suggest that CI downregulation is one adaptive trajectory ^15^ which is available to these SDH-deficient cells; however, since CI suppression is not a universal feature of SDH-deficient cancer models ^22,23^, we wondered whether other, CI-independent adaptive trajectories exist. To this end, we cultured five SDHB-KO clones in media containing pyruvate under standard conditions for several months. At each passage, we quantified total cell numbers and re-plated a known number of cells (roughly 10% of the population, to avoid bottlenecking), which allowed us to calculate average inter-passage proliferation rates for each clone. All five clones roughly doubled their average proliferation rates over a span of 80-120 cumulative population doublings (approximately 3-4 months in culture) (Figure 1G, Figure S1D-E). This proliferation rate increase was stereotyped among clones (Figure S1D-E) and reasonably well-described by a sigmoidal curve with an inflection point (the point at which proliferation rate begins to plateau) at around 40 cumulative population doublings (Figure S1F-G, Supplementary Table 1).

To determine whether this proliferation rate increase corresponded to a reduction in aspartate limitation, we thawed unadapted, or ‘early passage’ (EP) versions of clones 1-3 along with their adapted, or ‘late passage’ (LP) counterparts and measured their proliferation with or without aspartate supplementation. Consistent with our long-term proliferation rate measurements, LP clones proliferated significantly faster than their EP counterparts, and their proliferation was only modestly improved by aspartate supplementation, indicating that they had largely overcome aspartate limitation (Figure 1H).

Long-term culture of mammalian cells is known to permit ‘phenotypic drift’ and can impose selective pressures specific to the idiosyncrasies of repeated passaging ^25,26^. To determine whether the proliferation phenomenon we observed in SDHB-KO clones is a nonspecific consequence of long-term culture versus a specific aspartate-dependent adaptation, we cultured EP clones 1-3 for 6 months in media containing excess aspartate (Figure S1H). These ‘aspartate-reared’ (AR) clones maintained consistently high inter-passage proliferation rates during the entire culture duration (Figure S1I), during which time they would not be expected to encounter selective pressures related to aspartate availability. After culturing these cells for 80-120 cumulative population doublings, we then withdrew aspartate and asked whether these AR clones more closely resembled EP or LP clones—that is, whether exogenous aspartate was able to prevent their adaptation (Figure S1H).

Strikingly, aspartate withdrawal in AR clones 1-3 significantly reduced proliferation rates to levels comparable with their EP counterparts (Figure S1J). To further examine whether these proliferative changes accompany changes in cellular aspartate levels, we used LCMS to quantify relative aspartate abundance in EP, AR, and LP clones 1-3. This revealed that, while LP clones have roughly twice as much aspartate as their EP counterparts, aspartate-weaned AR clones had comparable or lower aspartate levels than their EP counterparts, suggesting that exogenous aspartate prevented their adaptation (Figure S1K).

Finally, we assessed whether the adapted cells suppressed CI expression by blotting for the CI subunit NDUFA8 in wild-type (WT) parental cells, KO clones 1-3, and a previously adapted SDHB-KO clone (PA) which had adapted by downregulating CI ^15^. Interestingly, while KO clone 1 showed reduced levels of NDUFA8 comparable to PA, clones 2 and 3 showed similar NDUFA8 expression to parental cells (Figure 1I). Blotting for another CI subunit, NDUFV2, largely replicated this finding (Figure S1L), prompting us to measure total CI activity in these cells using a modified respirometry protocol. All three LP clones had significantly lower basal oxygen consumption rates (OCR) than wild-type parental cells, consistent with SDH activity contributing to overall OCR ^15^, which we also confirmed by analyzing parental cells treated with the SDH inhibitor Atpenin A5 (AA5) (Figure 1J-K). Consistent with our western blot results, clone 1 showed slightly decreased OCR compared to AA5-treated WT cells, approaching the low OCR of PA and suggesting that this clone adapted by suppressing CI. However, the OCRs of clones 2 and 3 were significantly higher than PA and even slightly higher than AA5-treated WT cells (Figure 1J-K). The residual OCR of these clones was abolished with rotenone treatment (Figure 1J), indicating that these adapted clones had impaired SDH activity but intact CI activity.

Overall, these results reaffirm that SDH deficiency can cause aspartate limitation in cancer cells, but they also demonstrate that cancer cells can progressively adapt to overcome this aspartate limitation. Here, we find that this adaptation occurs with relatively slow kinetics and is not an inevitable or generalized consequence of long-term culture but instead appears to be a specific response to insufficient aspartate availability. These results also reveal that—while CI suppression is one adaptive trajectory available to SDH-deficient cells—CI loss is not required for adaptation. We subsequently sought to better understand the metabolic and functional features of the CI-independent adaptive trajectory.

### CI-dependent and CI-independent adaptation produces metabolically distinct outcomes

We focused on characterizing SDHB KO clone 2, which had adapted to overcome aspartate limitation without losing CI activity (Figure 1G-K, Figure S2K-L). To verify that genetic CI loss is sufficient to adapt this clone, we used CRISPR/Cas9 to knock out NDUFA8 in EP clone 2, generating a cell line which we refer to as ‘A8KO’ (Figure 2A-B). A8KO cells had virtually no mitochondrial oxygen consumption, reduced whole-cell NAD^+^/NADH, increased whole-cell aspartate levels, and reduced aspartate limitation compared to their parental EP clone 2 cells (Figure 2C-D, S2A-B), confirming that genetic CI loss is sufficient for adaptation in this clone.

**Fig. 2.**
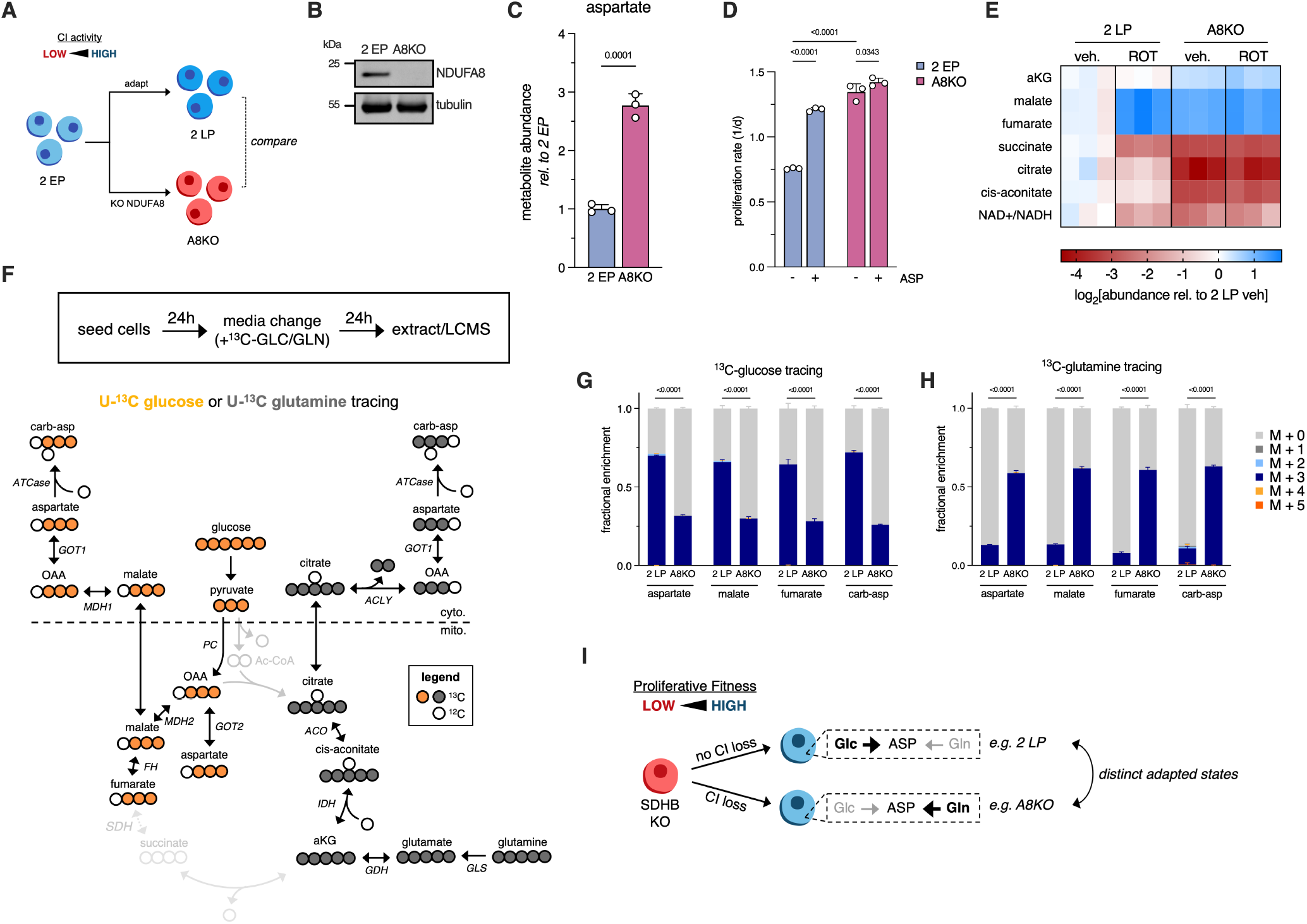
CI-high and CI-low adapted SDH-deficient cells are metabolically distinct. (**A**) Schematic illustrating the creation of NDUFA8-KO (A8KO) cells from EP SDHB KO clone 2 (2 EP), and the subsequent comparison between A8KO and LP SDHB KO clone 2 (2 LP) (which had adapted without suppressing CI). (**B**) Representative western blot showing levels of NDUFA8 and tubulin loading control in 2 EP and A8KO. (**C**) Relative whole-cell aspartate levels (mean +/-S.D.) in 2 EP and A8KO 24 hours after media change. Levels are normalized to 2 EP. (n=3) (**D**) Proliferation rates (mean +/-S.D.) of 2 EP and A8KO with or without 20 mM aspartate supplementation. (n=3) (**E**) Heatmap depicting log2-transformed relative abundances of the indicated metabolites in 2 LP and A8KO treated with vehicle control (veh.) or 50 nM rotenone (ROT) for 24 hours. Abundances are normalized to the vehicle-treated 2 LP condition. (n=3) (**F**) Schematic depicting U-^13^C glucose or U-^13^C glutamine tracing to determine routes of alternative aspartate synthesis in SDH-deficient cells. The experimental procedure for results in panels G-H is shown in the box. (**G**) Fractional isotopolog distributions for the indicated metabolites in 2 LP and A8KO following 24 hours of tracing in media containing U-^13^C glucose. P-values represent results from comparing the M+3 fractions. (n=3) (**H**) Fractional isotopolog distributions for the indicated metabolites in 2 LP and A8KO following 24 hours of tracing in media containing U-^13^C glutamine. P-values represent results from comparing the M+3 fractions. (n=3) (**I**) Schematic illustrating the existence of at least two distinct adaptive trajectories in SDHB-KO cells, which differ in complex I activity status and in the major metabolic source of aspartate [glucose (Glc) vs. glutamine (Gln)]. Statistical significance determined using an unpaired t-test (**C**), ordinary two-way ANOVA and uncorrected Fisher’s LSD with single pooled variance (**D**) or ordinary two-way ANOVA and Tukey’s multiple comparisons test with a single pooled variance (**G-H**).

LP clone 2 and A8KO arose from the same parental cell population but differ in their CI status (Figure 2A). Both show an ‘adapted’ proliferative phenotype (Figure 1H, 2D), but do they represent two distinct metabolic states, or convergence to the same adapted metabolic state? To answer this question, we used LCMS to profile the steady-state levels of a collec-tion of TCA cycle-relevant metabolites in these two cell lines. This revealed considerable metabolic differences, with A8KO showing higher levels of alpha-ketoglutarate (aKG), malate, and fumarate, and lower succinate, citrate, cis-aconitate, and NAD^+^/NADH than LP clone 2 (Figure 2E). Many of the metabolites that were less abundant in A8KO were down-stream of NAD^+^-dependent metabolic reactions, while many that were enriched in A8KO were downstream of NADH-dependent reactions (Figure S2C), suggesting mitochondrial redox as a likely driver of these metabolic differences. Consistent with this idea, rotenone treatment shifted the metabolite levels of LP clone 2 to resemble those of A8KO (but had little effect on A8KO, as these cells already have no CI activity) (Figure 2E).

These results suggest that A8KO adopts a reductive metabolic configuration, while LP clone 2 maintains some level of oxidative mitochondrial metabolism. To independently confirm this possibility, we cultured these cells in uniformly ^13^C-labeled glucose or glutamine for 24 hours and used LCMS to query labeling of cis-aconitate from these two sources. In SDH-deficient cells, oxidative reactions in the residual TCA cycle should produce cis-aconitate labeled M+2 and M+5 from U-^13^C glucose, while reductive TCA activity should generate M+5 labelled cis-aconitate from U-^13^C glutamine ^27^ (Figure S2D). Indeed, while LP clone 2 labeled roughly half of cis-aconitate pools M+2/M+5 from U-^13^C glucose, cis-aconitate labeling from glucose was virtually undetectable in A8KO (Figure S2E-F). Conversely, while a small fraction of cis-aconitate was labeled M+5 from U-^13^C glutamine in LP clone 2, a significantly larger proportion was labeled in A8KO (Figure S2G-H). Overall, this argues that the overall central carbon metabolic ‘states’ of LP clone 2 and A8KO are distinct, likely driven by CI-dependent changes in mitochondrial redox ^15,20,28^.

Do these two distinct adapted states synthesize aspartate using distinct pathways? To address this, we queried the afore-mentioned isotope tracing dataset for labeling of aspartate and nearby metabolites, which can in some cases distinguish between alternative aspartate synthesis pathways ^13–15^. In this case, aspartate synthesized via the RCQ pathway should be labeled M+3 from U-^13^C glutamine, while aspartate synthesized via PC should be labeled M+3 from U-^13^C glucose (Figure 2F). Indeed, this analysis demonstrated another stark metabolic contrast between LP clone 2 and A8KO: while the majority (70%) of aspartate was labeled M+3 from glucose in LP clone 2, the majority of aspartate in A8KO was unlabeled from glucose (Figure 2G, S2I). Instead, the majority of aspartate in A8KO was glutamine-derived (Figure 2H, S2M, with a smaller portion labeled from glucose (Figure 2G, S2I). These differential labeling patterns were mirrored in malate, fumarate, and carbamoyl-aspartate (carb-asp) (Figure 2G-H, S2J-L, N-P), which are generally in equilibrium with aspartate pools ^24,29^. Approximately 85-90% of aspartate was labeled either from U-^13^C glucose or glutamine in both cell lines (Figure 2G-H), demonstrating that the vast majority of aspartate in these cells is synthesized *de novo* and not scavenged from the environment or produced from other metabolic substrates.

We conclude that LP clone 2 and A8KO represent two fundamentally distinct adapted metabolic states derived from the same unadapted SDH-deficient clone. While these two adapted states have similarly high proliferative fitness, they differ both in their overall mitochondrial metabolic configuration (reductive vs oxidative) and in the specific alternative aspartate synthesis that facilitates the adaptation—SDH-deficient cells with suppressed CI synthesize a majority of aspartate from glutamine, while adapted cells that retain CI activity primarily employ glucose-dependent aspartate synthesis (Figure 2I).

### Differential GOT enzyme usage distinguishes adaptive trajectories of SDH-deficient cells

While isotope tracing indicated different metabolic sources of aspartate in these two adapted states, we next turned our attention to delineating the exact alternative aspartate synthesis pathways at play in these two contexts. We previously found that 143B cells with concurrent SDH and CI loss depend on mitochondrial pyruvate import via the mitochondrial pyruvate carrier (MPC)—both to provide substrate to pyruvate carboxylase (PC) and to facilitate aKG generation via glutamate pyruvate transaminase 2 (GPT2) ^15^ (Figure 3A). To determine whether SDH-deficient, CI-intact adapted cells also depend on mitochondrial pyruvate import, we generated MPC1 knockout and addback LP clone 2 and A8KO pools using CRISPR/Cas9 and lentivirus (Figure 3B), which were maintained in media containing excess aspartate. In both knockout lines, aspartate withdrawal induced proliferative defects which were absent or less severe in the corresponding MPC1 addbacks (Figure 3C-D. Aspartate levels were equivalently depleted in both MPC1 knockout lines compared to their addback counterparts (Figure 3E), confirming that MPC activity is necessary for aspartate synthesis in both adapted states.

**Fig. 3.**
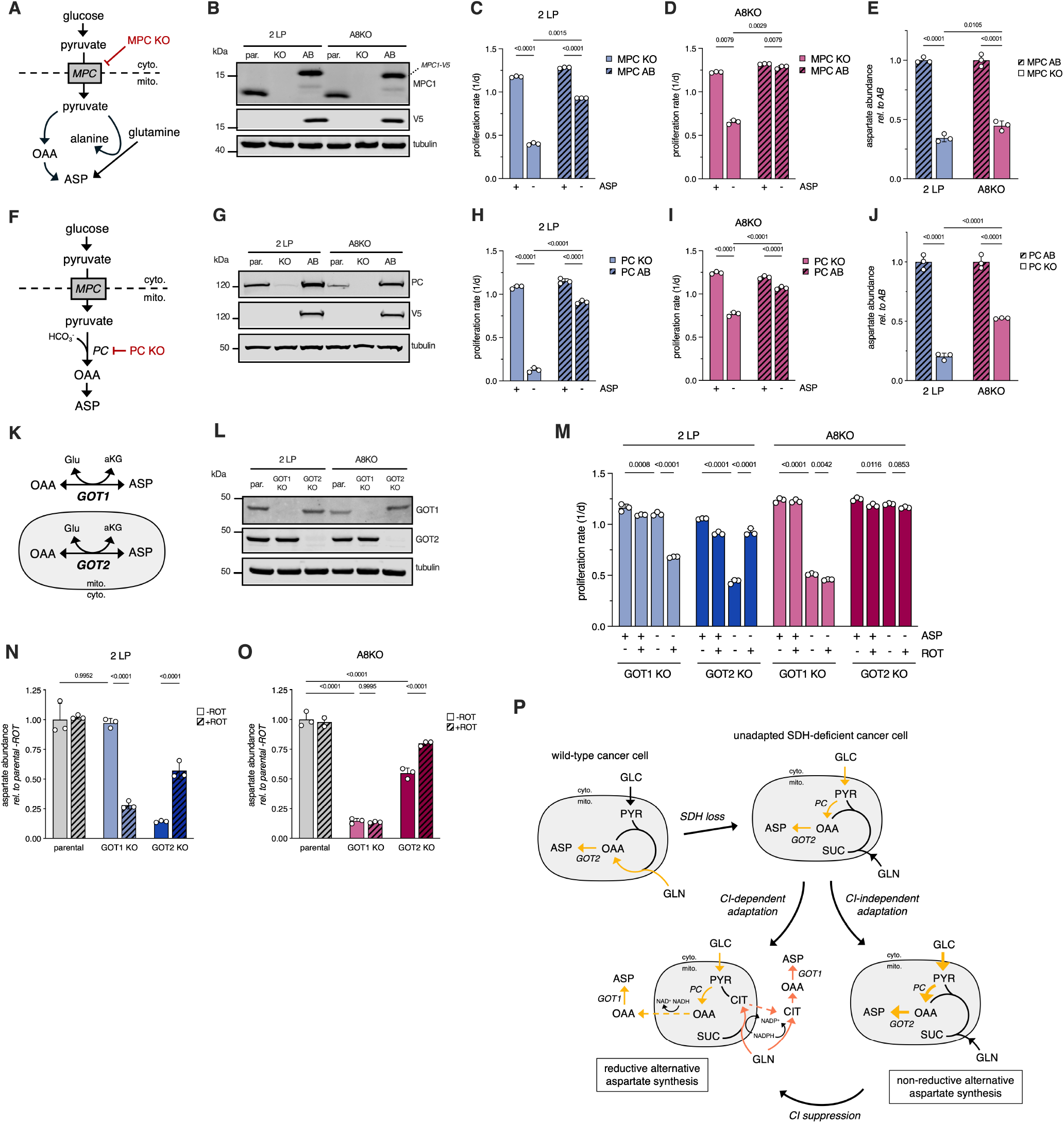
Shared and distinct aspartate-related metabolic dependencies of CI-high and CI-low adapted SDH-deficient cells. (**A**) Schematic illustrating the role of the mitochondrial pyruvate carrier (MPC) in alternative aspartate synthesis and its knockout using CRISPR/Cas9 (**B**) Representative western blot showing levels of MPC1, V5-tagged MPC1, and tubulin loading control in parental, MPC1-knockout (KO) and MPC1-addback (AB) cells for late passage SDHB-KO clone 2 (2 LP) and 2 EP NDUFA8-KO (A8KO). (**C-D**) Proliferation rates (mean +/-S.D.) of MPC KO and addback (AB) 2 LP (**C**) or A8KO (**D**) with or without 20 mM aspartate supplementation. (n=3) (**E**) Relative aspartate levels (mean +/-S.D.) of MPC KO and addback (AB) 2 LP and A8KO 24 hours after media change. Levels are normalized to each corresponding AB. (n=3) (**F**) Schematic illustrating the role of pyruvate carboxylase (PC) in alternative aspartate synthesis and its knockout using CRISPR/Cas9 (**G**) Representative western blot showing levels of PC, V5-tagged PC, and tubulin loading control in parental, PC-knockout (KO) and PC-addback (AB) 2 LP and A8KO cells (**H-I**) Proliferation rates (mean +/-S.D.) of PC KO and addback (AB) 2 LP (**H**) or A8KO (**I**) with or without 20 mM aspartate supplementation. (n=3) (**J**) Relative aspartate levels (mean +/-S.D.) of PC KO and addback (AB) 2 LP and A8KO 24 hours after media change. Levels are normalized to each corresponding AB. (n=3) (**K**) Schematic illustrating compartmentalized aspartate synthesis by GOT1/2 (**L**) Representative western blot showing levels of GOT1, GOT2, and tubulin loading control in parental, GOT1-KO and GOT2-KO 2 LP and A8KO cells (**M**) Proliferation rates (mean +/-S.D.) of GOT1- and GOT2-KO 2 LP and A8KO cells in all combinations of 20 mM aspartate and 50 nM rotenone treatments. (n=3) (**N-O**) Relative aspartate levels (mean +/-S.D.) of parental, GOT1-, and GOT2-KO 2 LP (**N**) or A8KO (**O**) cells 24 hours after treatment with vehicle control or 50 nM rotenone. Levels are normalized to each corresponding parental cell vehicle treatment. (n=3) (**P**) Schematic depicting aspartate synthesis in wild-type and SDH-deficient 143B cells before and after adaptation along two distinct trajectories. Abbreviations: GOT1, glutamic-oxaloacetic aminotransferase 1; GOT2, glutamic-oxaloacetic aminotransferase 2; ASP, aspartate; OAA, oxaloacetate; cyto., cytosol; mito., mitochondria; GLC, glucose; GLN, glutamine; PYR, pyruvate; SDH, succinate dehydrogenase; SUC, succinate; PC, pyruvate carboxylase; CI, respiratory complex I; CIT, citrate. Statistical significance determined using an ordinary two-way ANOVA and uncorrected Fisher’s LSD with single pooled variance (**C-E, H-J**) or ordinary two-way ANOVA and Tukey’s multiple comparisons test with a single pooled variance (**M-O**).

In agreement with these findings, the well-characterized MPC inhibitor UK-5099 caused aspartate-dependent proliferative defects in LP clones 1-3 and A8KO cells, but not wild-type parental 143B cells (Figure S3A-F). Altogether, these results indicate that mitochondrial pyruvate import is necessary for alternative aspartate synthesis and proliferation in both CI-suppressed and CI-intact adapted SDH-deficient cells.

Following import into mitochondria, pyruvate can be converted into the aspartate precursor oxaloacetate (OAA) by the enzyme pyruvate carboxylase (PC) (Figure 3F). We again used CRISPR/Cas9 and lentivirus to determine the requirement for PC in LP clone 2 and A8KO by generating PC-knockout and addback pools from both of these cell lines (Figure 3G). In both cell lines, PC loss led to an aspartate-dependent proliferative defect that was ameliorated in addbacks; however, the magnitude of this defect was larger in LP clone 2 than A8KO despite a nearly equivalent knockout efficiency (Figure 3H-I). Aspartate levels mirrored this trend, with PC loss causing a 75% aspartate reduction in LP clone 2 compared to a 50% reduction in A8KO (Figure 3J). Overall, these results indicate that PC is essential for aspartate synthesis and fitness in CI-intact adapted SDH-deficient cells, and important but slightly less essential in CI-suppressed adapted cells. This discrepancy fits nicely with the earlier isotope tracing results, which indicated that CI-intact cells (i.e. LP clone 2) synthesize a majority of aspartate from glucose, which requires PC activity, while CI-suppressed cells (i.e. A8KO) favor glutamine-dependent aspartate synthesis, which does not require PC activity (Figure 2E-G).

Finally, we turned our attention to the terminal step of alternative aspartate synthesis: the conversion of OAA into aspartate. This reaction is catalyzed by two paralogs of glutamic-oxaloacetic transaminase (GOT) enzymes present in most cells: cytosolic GOT1 or mitochondrial GOT2, which catalyze identical reactions in their respective cellular compartments (Figure 3K). Since the isotope tracing strategy employed previously (Figure 2E) is unable to distinguish between GOT1- and GOT2-synthesized aspartate, we used CRISPR/Cas9 to knock out GOT1/2 in LP clone 2 and A8KO (Figure 3L) to determine their relative contributions to aspartate pools and proliferative fitness.

In contrast to the previous MPC/PC knockout experiments, these experiments revealed a clear distinction in pathway usage between LP clone 2 and A8KO. While GOT1 knockout in LP clone 2 did not affect proliferation, GOT2 knockout in the same cell line significantly impaired proliferation in an aspartate-dependent manner (Figure 3M). The converse was true for A8KO, where GOT2 loss was completely tolerated while GOT1 loss was detrimental but fully rescuable by aspartate (Figure 3M).

Since CI inhibition was capable of changing the metabolic phenotype of LP clone 2 to resemble that of A8KO (Figure 2D), we tested whether rotenone treatment impacted GOT dependency in these cell lines. Strikingly, rotenone treatment ‘flipped’ the GOT dependency of LP clone 2 to match that of A8KO: rotenone inhibited the proliferation of GOT1 knock-out LP clone 2 (in an aspartate-dependent manner), whereas rotenone rescued the growth defect in GOT2 knockout LP clone 2 (Figure 3M). As expected, rotenone had no effect on proliferation in either GOT1 or GOT2 knockout A8KO cells (Figure 3M).

In these experiments, proliferative defects were considerable but not complete, despite nearly 100% knockout efficiencies (Figure 3L-M). Rather than reflecting mixed GOT1/2 usage by these cell lines, we suspected that this was an artifact of the experimental setup, in which GOT KO cells were cultured in aspartate-containing media, and 3-day proliferation assays were commenced by withdrawing aspartate in the relevant conditions. To test whether the incomplete proliferative defects reflected cells’ spending their residual aspartate pools following aspartate withdrawal, we determined the proliferation rates of these conditions between days 2 and 3 of the assay. Consistent with our hypothesis, this revealed that—after residual aspartate pools are spent—GOT2 knockout LP clone 2 or GOT1 knockout A8KO were virtually unable to proliferate without supplemental aspartate, as were rotenone-treated GOT1 knockout LP clone 2 cells (Figure S3G).

Cellular aspartate pools in these different cell lines and conditions largely mirrored the proliferation phenotypes: GOT1 knockout caused large (>75%) aspartate decreases in A8KO cells or rotenone-treated LP clone 2 but did not affect aspartate abundance in untreated LP clone 2 (Figure 3N-O). GOT2 knockout, meanwhile, caused similarly large aspartate depletion in LP clone 2, and this effect was partially rescuable by rotenone (Figure 3N). Interestingly, GOT2-KO A8KO cells showed moderately depleted aspartate pools, though this decrease was not enough to inhibit proliferation compared to parental cells (Figure 3M, O). Overall, these results suggest that adapted SDH-deficient cells depend on either GOT1 or GOT2 (but not both) for maximal aspartate synthesis and proliferation, and that CI activity enforces this largely binary phenotype.

Together with our previous study ^15^, these results allowed us to formulate a comprehensive model for adaptive aspartate synthesis in SDH-deficient 143B cells. While wild-type 143B cells synthesize most aspartate from glutamine via oxidative TCA cycling and GOT2 activity, acute SDH loss produces a low-fitness metabolic state in which a relatively small amount of aspartate is produced from glucose via PC and GOT2 (hereafter referred to as the ‘PC-GOT2 pathway’) (Figure 3P). These cells can overcome aspartate limitation by suppressing CI activity, which licenses GOT1-dependent alternative aspartate synthesis ^30^ via reductive glucose- and glutamine-derived pathways (Figure 2F-G, 3M, O-P). Alternatively, these cells can adapt via a distinct route that preserves CI activity and aspartate synthesis via the PC-GOT2 pathway. Curiously, SDH-deficient cells are not irreversibly ‘committed’ to the CI-intact adapted metabolic state, since pharmacological CI suppression reverts their metabolic state and dependencies to that of CI-suppressed SDH-deficient cells (Figure 2D, 3M-N, P).

### PC expression can be rate-limiting for aspartate synthesis and proliferation in SDH-deficient cells

We found above that SDH-deficient cells can adapt to overcome aspartate limitation via either of two distinct adaptive trajectories. While we previously demonstrated CI suppression as a mechanism by which SDH-deficient cells can adapt to engage reductive alternative aspartate synthesis pathways ^15^, we sought a mechanism by which cells can adapt via the CI-independent trajectory. Our previous data suggested that these cells adapted by increasing aspartate synthesis via the PC-GOT2 pathway (Figure 3P). To confirm this, we quantified the labeling of aspartate from U-^13^C glucose in EP and LP clone 2. Aspartate synthesized via the PC-GOT2 pathway should be labelled M+3 from labeled glucose (Figure 4A), and indeed, LP clone 2 demonstrates both higher abundance (Figure 4B) and fractional enrichment (Figure S4A) of M+3 aspartate from glucose than its EP counterpart, consistent with adaptive upregulation of PC-GOT2 pathway activity in this clone.

**Fig. 4.**
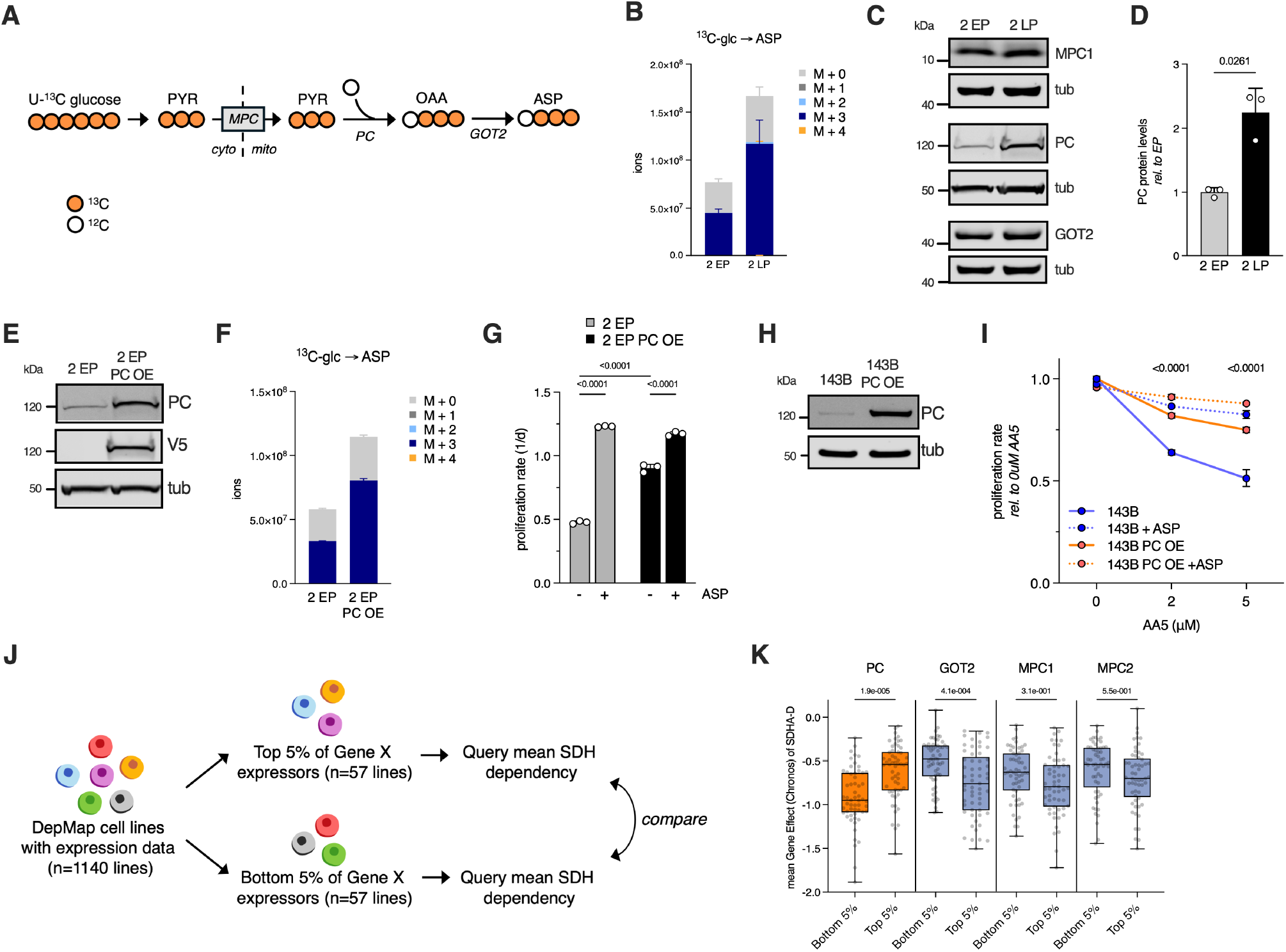
PC expression can be rate-limiting for aspartate synthesis and proliferation in SDH-deficient cells. (**A**) Schematic depicting U-^13^C glucose tracing to monitor aspartate synthesis via the MPC-PC-GOT2 pathway (**B**) Absolute isotopolog distributions (mean +/-S.D.) for aspartate in early passage (EP) or late passage (LP) SDHB-KO clone 2 following 24 hours of tracing in media containing U-^13^C glucose. (n=3) (**C**) Representative western blots showing levels of MPC1, PC, GOT2, and tubulin loading control in 2 EP/LP (**D**) Quantification of PC protein levels from 2 EP/LP using western blotting. Statistical significance determined using Welsh’s t-test. (n=3) (**E**) Representative western blots showing levels of PC, V5-tagged PC, and tubulin loading control in 2 EP or PC-overexpressing 2 EP (2 EP PC OE). (**F**) Absolute isotopolog distributions (mean +/-S.D.) for aspartate in 2 EP and 2 EP PC OE after 24 hours of tracing in media containing U-^13^C glucose. (n=3) (**G**) Proliferation rates (mean +/-S.D.) of 2 EP and 2 EP PC OE with or without 20 mM aspartate (ASP) supplementation. Statistical significance determined using an ordinary two-way ANOVA and uncorrected Fisher’s LSD with a single pooled variance. (n=3) (**H**) Representative western blots showing levels of PC and tubulin loading control in wild-type 143B cells or PC-overexpressing 143B cells (143B PC OE). (**I**) Proliferation rates (mean +/-S.D.) of 143B and 143B PC OE at the indicated doses of Atpenin A5 (AA5) with or without 20 mM aspartate (ASP) supplementation. Proliferation rates are normalized to the 0 *µ*M 5 treatment condition for each respective trace. P-values shown correspond to statistical testing comparing the relative proliferation rate of 143B vs. 143B PC OE at the indicated dose of AA5. (n=3) (**J**) Schematic illustrating the approach in (**K**), described further in the main text. (**K**) Mean SDHA-D DepMap gene effect scores for the top and bottom 5% of expressors of the indicated metabolic genes. Statistical significance determined using an ordinary one-way ANOVA and Sidak’s multiple comparisons test with a single pooled variance. (n=57) Statistical significance determined using Welch’s t-test (**D**), an ordinary two-way ANOVA and uncorrected Fisher’s LSD with single pooled variance (**G**), ordinary two-way ANOVA and Tukey’s multiple comparisons test with a single pooled variance (**I**), or ordinary one-way ANOVA with Sidak’s multiple comparisons test with a single pooled variance (**K**).

To determine whether this clone may have adapted by up-regulating one of the enzymes in the PC-GOT2 pathway, we quantified the abundance of MPC1, PC, and GOT2 in EP and LP clone 2. While unadapted and adapted cells had roughly equal amounts of MPC1 and GOT2, adapted cells showed a >2-fold increase in protein levels of PC (Figure 4C-D), suggesting that PC expression is rate-limiting for aspartate synthesis. To test this possibility, we used lentivirus to overexpress V5-tagged PC in EP clone 2 (Figure 4E) and measured effects on aspartate synthesis and proliferation. Strikingly, PC overexpression in this clone increased aspartate levels, with labeling patterns consistent with increased aspartate synthesis via the PC-GOT2 pathway (Figure 4F, S4B). PC overexpression accordingly increased proliferation and decreased aspartate limitation, producing cells that were functionally similar to LP clone 2 (Figure 4G, 1H). To rule out that PC overexpression caused these effects by decreasing CI activity, we quantified whole-cell NAD^+^/NADH and examined the levels and labeling of succinate, whose synthesis in SDH-deficient cells is highly responsive to CI activity ^24^ (Figure 2D). Compared to EP clone 2, PC-overexpressing EP clone 2 had no significant change in NAD^+^/NADH or succinate levels, with succinate labeling consistent with synthesis from glutamine (Figure S4C-F), indicating that PC overexpression does not appreciably alter CI activity in this context.

To check whether this adaptive mechanism generalized beyond this single clone, we blotted for MPC1, PC, and GOT2 in EP/LP clone 3, which had also preserved CI activity during adaptation (Figure 1K). Similar to clone 2, LP clone 3 showed a specific upregulation of PC compared to its EP counterpart (Figure S4G-H), suggesting that PC upregulation is a convergent feature of CI-independent adaptation to SDH deficiency. We next sought an orthogonal method of testing whether PC expression alone determined sensitivity to SDH deficiency in 143B cells. We reasoned that, if this were true, then overexpressing PC in wild type 143B cells should desensitize them to the antiproliferative effects of pharmacological SDH inhibition. Indeed, PC-overexpressing cells were significantly desensitized to the SDH inhibitor Atpenin A5 (AA5) compared to wild-type parental controls (Figure 4H-I).

Finally, we sought to generalize these results to a larger collection of models. To do so, we queried DepMap ^31^, which integrates expression profiles for over one thousand cancer cell lines with cell-line specific gene dependencies, and leveraged the fact that DepMap cell lines show a relatively wide range of sensitivity to knockout of SDH subunits (Figure S4I). We reasoned that—if alternative aspartate synthesis and proliferation were indeed generally limited by PC protein levels—then cell lines with higher basal PC expression should tend to show decreased SDH dependency. Thus, we averaged the gene effect scores for all four SDH subunits (SDHA-D) as a proxy for ‘SDH dependency’ and correlated this with relative PC mRNA expression across 1140 DepMap cell lines, which returned a modest but significantly non-zero correlation (Figure S4J).

While this correlation was weak, it’s important to note that—beyond potentially reflecting varying SDH knockout ef-ficiencies—this dataset represents many different tissue types, growth habits, and dozens of media formulations that differ in components (pyruvate, salvageable nucleotide precursors, amino acids) known to influence SDH dependency in some cases ^15,24,32^. Nevertheless, we did not observe a similar correlation between SDH dependency and expression of other PC-GOT2 pathway enzymes (MPC1/2, GOT2) (Figure S4K-M), or other relevant metabolic nodes including GOT1, MDH1/2, and PDHA1—a component of the pyruvate dehydrogenase complex which competes with PC for mitochondrial pyruvate as substrate (Figure S4N-Q).

Reasoning that perhaps a relationship between PC expression and SDH dependency wouldn’t be strictly linear (for example, if there were a certain threshold of PC expression below which aspartate synthesis constrained cell growth or viability) we performed a modified analysis in which we extracted the cell lines with the top and bottom 5% of PC expression and compared the mean SDH gene effect between these two populations (each of which consisted of 57 cell lines) (Figure 4J). This analysis revealed that cell lines in the bottom 5% of PC expression showed significantly larger SDH gene effect scores than those in the top 5% (Figure 4K). Replicating the analysis on GOT2, MPC1/2, GOT1, MDH1/2, and PDHA1 again suggested that this effect was specific to PC expression and not a more general consequence of cells with high metabolic gene expression (Figure 4K, S4R). Overall, these results indicate that PC expression can be rate-limiting for aspartate synthesis and proliferation of SDH-deficient cells and nominate PC upregulation as a mechanism by which SDH-deficient cells can adapt via the CI-independent trajectory (Figure 3P).

### SDH-deficient cells adapt to overcome pyrimidine insufficiency and replication stress

Having defined two distinct adaptive trajectories through which SDH-deficient cells can overcome aspartate-dependent proliferative defects, we sought to understand the broader metabolic and functional consequences of adaptation. To gain a more complete view of adaptive changes to aspartate metabolism, we used LCMS to quantify the abundance of several aspartate fates (Figure 5A) in EP, AR, and LP clones 1-3. In particular, we recently reported that SDH loss disproportionately impairs the consumption of aspartate into *de novo* pyrimidine synthesis due to a succinate-mediated competitive inhibition of the first enzymatic step in this pathway ^24^. Consistent with the idea that pyrimidine synthesis impairment is a specific metabolic hurdle for adapting SDH-KO cells to overcome, levels of the pyrimidine intermediates carbamoyl-aspartate and orotate and product UMP were significantly higher in all three LP clones compared to their EP versions (Figure 5B-D). Importantly, levels of these metabolites were comparable between EP and AR clones (Figure 5B-D), once again suggesting that these metabolic changes occur in response to aspartate limitation and not extended culture.

**Fig. 5.**
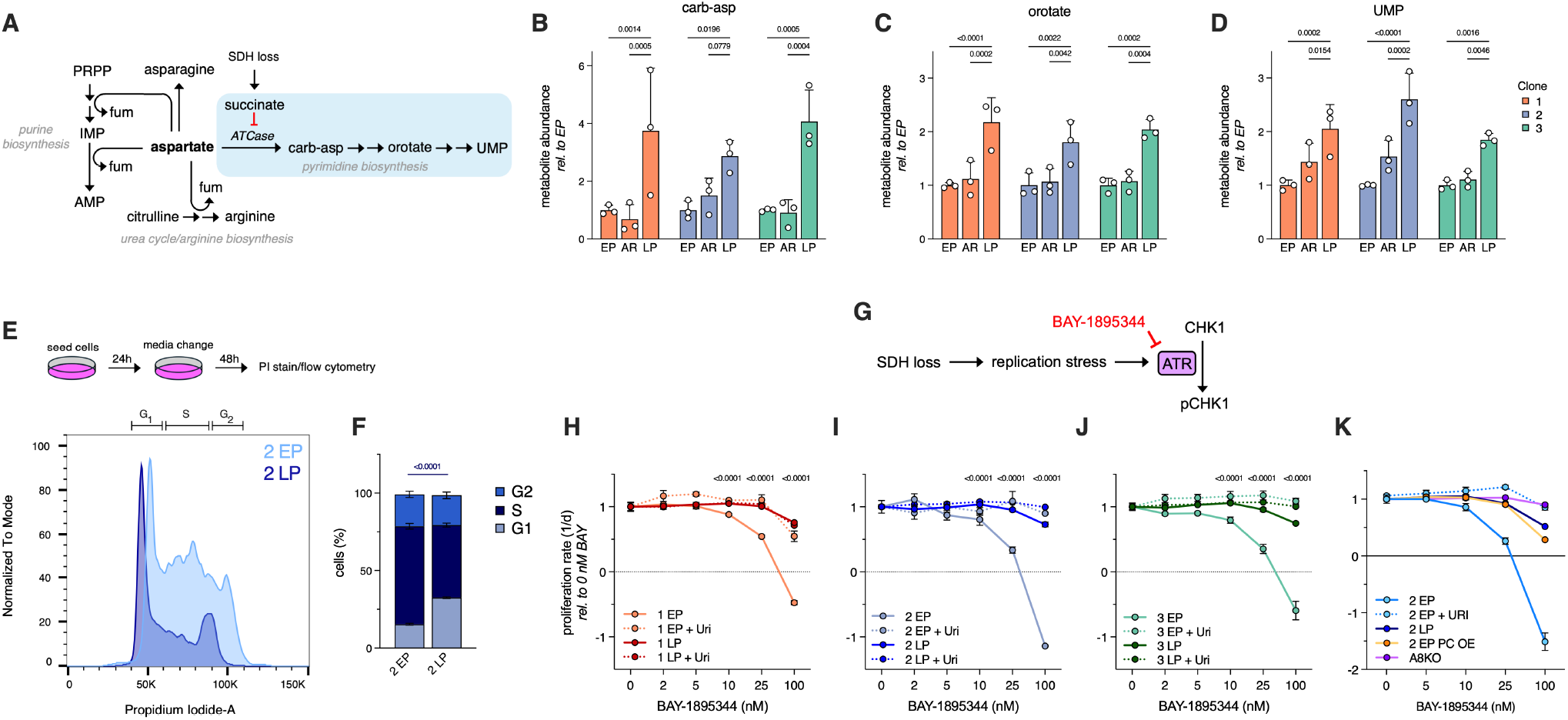
SDH-deficient cells adapt to overcome pyrimidine deficiency and replication stress. (**A**) Schematic depicting metabolic fates of aspartate in proliferating cells and the effect of SDH loss on pyrimidine biosynthesis ^24^ (**B-D**) Relative levels of carbamoyl-aspartate (carb-asp) (**B**), orotate (**C**), and uridine monophosphate (UMP) (**D**) (mean +/-S.D.) in early passage (EP), aspartate reared (AR), and late passage (LP) SDHB knockout clones 1-3. Levels are normalized to each corresponding EP clone. (n=3) (**E**) Schematic illustrating the cell cycle analysis experimental setup and normalized univariate histograms of propidium iodide (PI) staining in early passage (EP) and late passage (LP) knockout clone 2. Approximate regions corresponding to *G*_1_, *S*, and *G*_2_ cell cycle phases are labeled at the top of the chart. One replicate of each cell line is shown for clarity. (**F**) Cell cycle phase distributions of 2 EP/LP quantified from data in (**E**). P-value shown corresponds to statistical testing comparing the fraction of cells in S phase in each condition. (n=3) (**G**) Schematic illustrating the finding that SDH loss causes replication stress, the role of ATR kinase in the replication stress response, and the ATR inhibitor BAY-1895344 (BAY). (**H-K**) Proliferation rates (mean +/-S.D.) of EP/LP SDHB knockout clones 1-3 (**H-J**) or 2 EP, PC-overexpressing 2 EP (2 EP PC OE), 2 LP, and NDUFA8-KO 2 EP (A8KO) (**K**) in the indicated doses of BAY, with or without 500 *µ*M uridine supplementation. Proliferate rates are normalized to the 0 nM BAY dose in each condition. P-values shown correspond to statistical testing comparing the relative proliferation rate of EP vs. LP at the indicated dose of BAY. Statistical significance determined using ordinary two-way ANOVAs and uncorrected Fisher’s LSD with single pooled variance.

Examining other metabolic fates of aspartate, we found that the AMP/IMP ratio—an indicator of aspartate-dependent purine synthesis ^29^—is also significantly or trending higher in LP clones compared to EP (Figure S5A-C), as are levels of argininosuccinate, a urea cycle intermediate synthesized in part from aspartate ^33^ (Figure S5D). Interestingly, levels of the amino acid asparagine were not higher in LP clones and were consistently low in AR clones (Figure S5E), suggesting potential feedback regulation of asparagine synthetase (ASNS) in response to aspartate supplementation. Overall, these results reveal that adapted SDH-deficient cells have broadly improved metabolic signatures of aspartate metabolism in line with their increased proliferative fitness.

One functional consequence of the pyrimidine synthesis impairment following SDH loss is pyrimidine deficiency and subsequent slowing of S-phase ^24^. To query whether this emer-gent phenotype changes during adaptation of SDH-deficient cells, we used PI staining and flow cytometry to quantify the cell cycle phase distribution of EP and LP SDHB KO clone 2 (Figure 5E). Consistent with our previous results, EP clone 2 had an enlarged fraction of cells in S-phase (62%) compared to parental 143B cells (generally 40-50% ^24^). The S-phase fraction decreased significantly in LP clone 2 (to 45%), consistent with these cells having overcome pyrimidine deficiency and the associated cell cycling defects (Figure 5E-F).

While SDH loss causes partial S-phase arrest, persistent cell growth in the absence of division leads to a pyrimidine-dependent increase in cell volume ^24^. Compared to their EP counterparts, all three LP clones had significantly reduced cell volumes (Figure S5F). Notably, treatment of EP/LP clone 2 with either aspartate or uridine (a salvageable pyrimidine precursor ^24^) reliably decreased cell volume (Figure S5G), demonstrating that this cell enlargement is attributable to aspartate limitation/pyrimidine deficiency.

S-phase stalling following pyrimidine deficiency is mediated by signaling pathways that collectively orchestrate the mitigation of DNA replication stress ^34^. One key player in this ‘replication stress’ response pathway is Ataxia Telangiectasia and Rad3-related (ATR), a kinase which senses DNA damage and phosphorylates the downstream CHK1 kinase (Figure 5G) ^35^. Consistent with SDH loss causing replication stress and ATR’s role as a critical replication stress response element, SDH loss sensitizes cancer cells to ATR inhibition ^24^. To test whether the apparent amelioration of pyrimidine deficiency and cell cycling defects in adapted cells corresponds to a decreased vulnerability to ATR inhibition, we measured the proliferation of EP and LP clones 1-3 in a titration of the ATR inhibitor BAY-1895344. Remarkably, all three LP clones were significantly less sensitive to BAY-1895344 than their EP counterparts, though they remained sensitive to doses above 100 nM (Figure 5H-J). Uridine supplementation rescued sensitivity in nearly all cases, confirming this phenotype’s dependence on pyrimidine deficiency and indicating that the drug is on-target (Figure 5H-J). Finally, we tested whether adapting EP clone 2 by either of the two trajectories we’ve described—by genetically silencing CI activity or overexpressing PC—is sufficient to ameliorate ATRi sensitivity. Strikingly, A8KO cells and PC-overexpressing EP clone 2 both had significantly reduced sensitivity to BAY-1895344, indistinguishable from LP clone 2 (Figure 5K).

Thus, in addition to adapting to increase their proliferative fitness and levels of aspartate and downstream fates, SDH-deficient cancer cells are capable of overcoming functional deficiencies associated with SDH loss—namely, the sensitivity to drugs targeting replication stress response pathways—via at least two distinct adaptive trajectories.

### Adaptation in SDH-deficient human pheochromocytoma cells

Finally, we sought to generalize our findings beyond 143B cells into a more physiologically relevant context. To do so, we used hPheo1 cells, which were derived from a human pheochromocytoma tumor ^36^ (one of the tumor types in which SDH acts as a tumor suppressor ^16^) and asked whether these cells were sensitive to SDH loss. Indeed, SDH inhibition in wild-type hPheo1 cells caused robust fumarate depletion, succinate accumulation, and depletion of aspartate and several of its fates, including pyrimidine precursors and argininosuccinate (Figure S6A).

Accordingly, in media containing pyruvate, SDH inhibition reduced hPheo1 proliferation, and this effect could be rescued with either aspartate supplementation or rotenone treatment (Figure S6B). These results indicate that SDH impairment induces aspartate limitation in this system, which can be overcome by direct aspartate replenishment or by CI suppression. SDH inhibition also increased mean per-cell volume, a phenotype which was rescued with aspartate or rotenone, suggesting that SDH loss also promotes replication stress in hPheo1 cells.

Next, we used CRISPR/Cas9 to generate three hPheo1 SDHB knockout clones (Figure S6D) and profiled their proliferation in long-term culture. Consistent with our pharmacological SDH inhibition results, all three KO clones initially proliferated at rates roughly half that of wild-type parental cells (Figure S6B, E-G). To varying degrees, all three clones progressively improved their proliferation rates over approximately 120 cumulative population doublings (3 months) (Figure S6E-G), suggesting that they were also capable of adapting to SDH loss. We functionally characterized clone 1 and verified that it indeed adapted to overcome aspartate limitation (Figure S6H). Interestingly, while aspartate levels were largely unchanged in late passage (LP) clone 1 compared to its early passage (EP) counterpart, LP cells had significantly higher levels of the pyrimidine intermediates carbamoyl-aspartate, orotate, and UMP, as well as higher levels of argininosuccinate (Figure S6I), consistent with increased aspartate metabolism into its fates.

Compared to its EP counterpart, LP KO clone 1 also showed a significantly reduced NAD^+^/NADH ratio and reduced succinate levels (Figure S6J), metabolic phenotypes associated with adaptive CI suppression in 143B cells (Figure 2D). To evaluate this possibility, we used respirometry to measure total mitochondrial oxygen consumption rates (OCR) in this clone and found that LP clone 1 had roughly 50% reduced mitochondrial OCR compared to EP clone 1 (Figure S6K), consistent with CI suppression. These data suggested that this KO clone had taken the CI-dependent adaptive route instead of the CI-independent route—consistent with this idea, clone 1 did not upregulate PC expression during adaptation, although both EP/LP clone 1 showed slightly more PC expression than wild-type parental cells (Figure S6L-M). Finally, we tested whether adaptation altered sensitivity to ATR inhibition and found that LP clone 1 was less sensitive to BAY-1895344 than EP clone 1 (Figure S6N).

Overall, these results suggest that adaptation to overcome aspartate limitation following SDH loss is a general capability of cancer cells, including cells derived from tissues relevant to SDH-associated tumorigenesis. These results are also consistent with a model whereby the CI-dependent and CI-independent adaptive routes are distinct and differentially engaged.

## Discussion

One way to reconcile the apparent paradox that SDH is a tumor suppressor is by invoking that cancer cells are able to adaptively overcome the liabilities associated with SDH loss. Here, we rigorously characterize the *in vitro* adaptation of SDH-deficient cancer cells and discover that adaptation can occur via at least two distinct mechanisms. These distinct mechanisms represent specific responses to limited availability of the amino acid aspartate and result in distinct adapted metabolic states. More concretely, SDH-deficient cells can either suppress CI activity, which modulates cellular redox state to license cytosolic alternative aspartate synthesis via GOT1, or they can preserve CI activity and instead upregulate PC, which promotes mitochondrial alternative aspartate synthesis via GOT2.

Beyond increasing aspartate abundance and proliferative fitness, either of these trajectories allow SDH-deficient cells to partially overcome pyrimidine deficiency, cell cycling defects, and sensitivity to a drug which we previously identified as a potential therapeutic vulnerability of SDH-deficient cancer cells ^24^, underscoring a need to better understand these adaptations with the eventual goal of treating SDH-associated malignancies. On a more basic level, in characterizing the CI-independent adapted state, we reveal a metabolic configuration where cancer cells differentially engage the electron transport chain (ETC) to synthesize the biosynthetic precursors needed for robust proliferation in the complete absence of oxidative TCA cycle activity.

While aspartate limitation is a general feature of SDH-deficient cancer cells ^13–15^, previous studies have presented mixed results concerning the pathways by which these cells circumvent or address this limitation. Some studies report a prominent role for glucose-derived aspartate synthesis via PC ^13,14^, but one of these studies found variability in the extent of PC induction by SDH-deficient tumors ^13^, while both suggested a potential contribution of glutamine-derived reductive TCA cycling ^13,14^. We and others have identified that both of these pathways are associated with and/or promoted by the redox consequences of CI suppression ^15,21^, which is consistent with findings of reduced CI activity in SDH-deficient cells and tumors ^13,15,37^ but at odds with reports of SDH-deficient cells with intact CI activity ^22,23^.

The present finding that aspartate limitation can be overcome with or without CI suppression consolidates these mixed results and suggests a unified framework highlighting the plasticity of adaptive aspartate synthesis (Figure 3P). Our results suggest that both adapted states (CI-suppressed and CI-intact) can be obtained from the same parent cell population, so determining whether the different phenotypes in the aforementioned SDH-deficient models are a result of stochastic or tissue-specific variables is an important goal for future work.

While previous studies have shown that SDH-deficient cancer cells and tumors can upregulate PC to facilitate alternative aspartate synthesis ^13,14^, we extend these findings by showing that PC expression can be rate-limiting for alternative aspartate synthesis even in SDH-deficient cells, and that basal PC expression can influence SDH dependency across cell lines. Given the fact that aspartate limitation in SDH-deficient cells can be overcome with either PC upregulation or CI downregulation, it’s tempting to speculate that these requirements contribute to the strict tissue tropism of SDH-deficient tumors ^16^—that is, that SDH-deficient tumors may preferentially arise in tissues or cells that have high PC expression or low CI activity, or are otherwise primed to maximally enact these changes following SDH loss. This hypothesis awaits experimental validation, however.

Along with our previous work ^15^, these results also highlight the fact that SDH-independent aspartate synthesis actually comprises three partially-overlapping pathways (glucose-derived and GOT1-dependent, glucose-derived and GOT2-dependent, and glutamine-derived and GOT1-dependent), which differ in their subcellular compartmentalization and metabolic dependencies (Figure 3P). While GOT1/2 are often discussed in the context of their roles in the malate-aspartate shuttle (MAS) ^38,39^, our finding that loss of the non-aspartate synthesizing paralog in each respective context (CI-intact/suppressed) did not impact proliferation indicates that their major functional roles in SDH-deficient cells are that of aspartate synthesis and not electron shuttling. Our finding that CI activity gates the largely binary choice of GOT1-versus GOT2-dependent aspartate synthesis in SDH-deficient cells is supported by studies which show redox-dependent control of GOT/MDH enzyme directionality in other contexts ^15,30,40^.

One motivation behind understanding cancer cell adaptation is that adaptation may phenotypically separate cancer cells from their non-cancerous counterparts and thus provide a therapeutic window ^11^. Indeed, while previous studies suggested PC as a potential target in SDH-deficient cancer cells ^13,14^, our results indicate that SDH-deficient cells which have adapted via the CI-dependent route are less reliant on PC activity than CI-intact SDH deficient cells. Considering this previously unappreciated plasticity, we would argue that prioritizing targets that are shared between adapted states—for example, MPC—may have a higher likelihood of success while still retaining selectivity for SDH-deficient cancer cells (Figure S3F).

Beyond SDH-mutant tumors, the alternative aspartate synthesis pathways investigated here may be differentially engaged and functionally relevant in other contexts, including diverse cancers without SDH mutations but with functional SDH deficiency ^41–44^, glutamine limitation (under which PC becomes essential for aspartate synthesis ^45^), hypoxia (during which both GOT1 and GOT2 have been implicated in supporting aspartate synthesis ^46,47^), *in vivo* growth of KRAS-mutant pancreatic cancer xenografts (which show surprising plasticity in GOT usage ^46,48^), and some non-small cell lung tumors and breast cancer metastases (both of which show increased PC-dependent anapleurosis ^49–51^).

While this study leaves many open questions—the mechanisms behind adaptive CI suppression or PC upregulation in SDH-deficient cells, the determinants behind the choice of adaptive trajectory, and the extent to which SDH-deficient cancer cells engage these alternative aspartate synthesis pathways *in vivo*—defining the scope of potential metabolic adaptations to SDH-dependent aspartate limitation will serve as a foundation from which to ask these more targeted questions once more robust models of SDH deficient tumorigenesis become available. Finally, we hope that this experimental system of cancer cell evolution in which the selective pressure (i.e. aspartate availability) is well-defined may find use as a model to study more general and far-reaching concepts of cancer metabolic adaptation.

## Methods

### Cell culture

143B cells were obtained from ATCC, authenticated using small tandem repeat (STR) profiling (ATCC), and monitored at least twice yearly for mycoplasma contamination (MycoProbe, R&D Systems). hPheo1 cells were received as a gift from Dr. Jerry Shaw (UT Southwestern). Cells were maintained in a humidified incubator at 37C and 5% CO2. Unless otherwise noted, cells were cultured in Dulbecco’s Modified Eagle’s Medium (DMEM) with 4.5g/L glucose and L-glutamine (Corning, 50-013-PC) supplemented with 3.7g/L sodium bicarbonate (Sigma, S5761), 1% penicillin/streptomycin (Gibco, 15140), 10% fetal bovine serum (Cytiva Hyclone), and 1mM sodium pyruvate (Sigma, P8574).

### Generation of knockout cell lines

Knockout cell lines were generated similar to what is previously described ^24^. sgRNAs targeting the genes of interest were purchased from Synthego—see the Key Resources Table for specific guide sequences. To generate a knockout, each sgRNA was resuspended in TE buffer at 100 *µ*M, then 1.47 *µ*L of each stock sgRNA was combined with 13.27 *µ*L SF buffer with supplement 1 (Lonza, V4XC-2032) and 4.33 *µ*L stock sNLS-spCas9 (Aldevron, 9212). The resulting mixture was incubated on the bench for 20 min., before being used to resuspend 100-200K 143B cells and electroporated using a 4D-Nucleofector (Amaxa, Lonza) using electroporation program FP-133. Nucleofected cells were then moved to a 12-well plate with pre-warmed media and expanded/assayed as necessary. After an SDHB-KO pool of 143B cells was generated, clones were obtained by limiting dilution plating (0.33 cells/well) in a 96-well plate. For ND-UFA8/MPC1/PC/GOT1/GOT2, the CRISPR pool was used for subsequent experiments. Gene KOs were confirmed using western blotting of both the CRISPR pools and each single-cell clone used in this study.

### Cell proliferation/volume assays

12-24 hours prior to starting proliferation assays, cells were seeded in 24-well plates at a density of 15-30K cells/well. At the start of each assay, cells were switched into fresh media (1ml/well) containing the indicated treatments, and three wells were washed once with DPBS (Gibco, 14190), detached with 1X 0.25% Trypsin-EDTA (Gibco, 25200), and counted using a Beckman Coulter Multisizer 4 instrument to establish an initial (‘time zero, *T*_0_’) cell count. For experiments involving the removal of aspartate or other components from the seeding media, all wells were washed once with DPBS before treatment media was added. 2-4 days after treatment, cells were washed, trypsinized, and counted as above, and average proliferation rates were determined according to the equation:

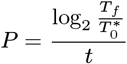

Where *P* = proliferation rate (population doublings/day), *T*_*f*_ = final cell count, 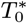 = initial cell count, averaged over three technical replicates, and *t* = time elapsed since *T*_0_ count in days. Per-cell volume data is automatically collected during counts, and mean per-cell volumes were extracted and plotted where appropriate. Unless otherwise noted, proliferation assays were conducted in media containing 1mM sodium pyruvate (Sigma, P8574). All proliferation rates were obtained in technical triplicate, and published results are representative of at least two independent experiments.

### SDHB knockout cell adaptation

For long-term adaptation, SDHB-knockout cells were passaged in 30cm culture dishes containing 35-40mL DMEM with sodium pyruvate (as described in ‘Cell Culture’). Cells adapted in media with 20mM aspartate were passaged in 10cm culture dishes containing 12mL media. At or near confluency, cells were washed, trypsinized, and counted (as in ‘Cell Proliferation Assays’), and 1-2 million cells (for -asp conditions) or 250-500,000 cells (for +asp conditions) were seeded into a new dish containing fresh media. A typical 30cm dish of confluent 143B cells holds 7-12 million cells, while a 10cm dish holds 4-7 million cells, so 10% of the cell population is carried over at passaging. Average proliferation rates between passages were estimated using the formula in ‘Cell Proliferation Assays,’ with 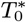 taken to be the number of cells seeded, *T*_*f*_ the final total cell count, and *t* the time elapsed in days between the two cell counts.

### Lentivirus production and infection

Expression constructs for SDHB, MPC1, and PC were obtained from the DNASU Plasmid Repository (Key Resources Table). Lentivirus was generated by transfection of HEK293T pLentiX cells with expression construct plasmid DNA and pMDLg/pRRE (Addgene, 12251), pRSV-Rev, (Addgene, 12253), and pMD2.G (Addgene, 12259) packaging plasmids using FuGENE transfection reagent (Fisher, PRE2693) in DMEM without FBS or penicillin-streptomycin. The supernatant containing lentiviral particles was filtered through a 0.45 *µ*M membrane (Fisher, 9720514) and stored at −80C before using. For infection, cells were plated at 20-50% confluence in multiwell plates, and the next day, media was changed to DMEM with 1 mM pyruvate supplemented with 8 µg/µL polybrene (Sigma, TR-1003-G) and a titration of viral supernatant. 24 hours later, media was again switched into selection media (DMEM containing 1 µg/mL blasticidin (Fisher, R21001)) until a parallel uninfected well of cells showed complete cell death (typically, 24-48 hours).

### Oxygen consumption measurements

For oxygen consumption measurements, cells were plated in Agilent Seahorse XFe96/XF Pro 96-well Cell Culture Microplates (Agilent, 103794) at densities from 30K-60K cells/well, which was optimized empirically for each cell line, and incubated in standard culture conditions overnight. The Seahorse sensor cartridge was equilibrated overnight in XF calibrant (Agilent, 100840) in a benchtop incubator at 37C. Approximately 18 hours later, cells were washed once with 180*µ*l/well Seahorse media (DMEM with 4.5g/L glucose and L-glutamine (Corning, 50-013-PC) and 1% penicillin/streptomycin (Gibco, 15140)), switched into 180*µ*l/well of Seahorse media with or without 5*µ*M Atpenin A5 (AA5), and incubated in a benchtop (non-CO2) incubator set to 37C for one hour. Separate 5*µ*M solutions of rotenone and antimycin A were prepared in Seahorse media and loaded into the first two ports of each well of a XFe96/XF Pro Extracellular Flux Assay Cartridge (Agilent, 103793). Final concentrations of rotenone and antimycin A upon injection into wells was 0.5*µ*M. Respirometry was performed on an Agilent Seahorse XFe96 Analyzer with the following protocol: equilibrate, 3 cycles of baseline measurement (mix 3 min., measure 3 min.), inject rotenone, 3 mix-measure cycles, inject antimycin, 3 mix-measure cycles. Following measurement, wells were gently washed once with DPBS, 10*µ*l/well protein lysis buffer (see ‘Western Blotting’) was added, and plates were freeze-thawed three times. Total protein concentrations per well were determined using a Pierce BCA Protein Assay Kit (Thermo Scientific, 23225) using a bovine serum albumin (BSA) standard curve prepared and measured in the leftover Seahorse Utility Plate. All respirometry measurements were normalized to the total protein amount per well, each well constitutes a technical replicate, and published results are rep-resentative of two independent respirometry experiments. For respirometry experiment in Figure S6, analysis was conducted on an Agilent Seahorse XFp instrument following similar protocols for sample preparation and running as described above. At the completion of this assay, cells were counted on a Beckman Coulter Multisizer 4 instrument, and oxygen consumption measurements were normalized to cell count rather than total protein.

### Protein Quantification via SDS-PAGE and Western Blotting

Protein lysis buffer—consisting of 1X RIPA (Millipore, 20-188), 5mM EDTA pH 8.0 (Sigma, 798681), and 1X Halt Protease Inhibitor Cocktail (Thermo Scientific, 1861279) in ultrapure water—was prepared, aliquoted, and stored at −20C. Adherent cells were either washed with DPBS and scraped in protein lysis buffer directly on the culture dish or else trypsinized, pelleted, washed once with 1X DPBS, and resuspended in protein lysis buffer. Samples were lysed on ice for 15 min with occasional vortexing, then spun at 17,000g for 10 minutes in a refrigerated centrifuge set to 4C to pellet insoluble components. Total protein concentrations were determined using a Pierce BCA Protein Assay Kit (Thermo Scientific, 23225), and solutions of equal protein concentration were prepared in 1X Bolt LDS Sample Buffer (Thermo Fisher, B0007) with 1X Bolt Sample Reducing Agent (Thermo Fisher, B0009). Samples were heated at 95C for 1 min, spun at 10,000g for 2 min at room temperature, and equal volumes were loaded onto precast 4-12% Bis-Tris Plus gels (Thermo Fisher, NW04125). SDS-PAGE was performed in 1X MES buffer (Thermo Fisher, B000202) at constant 180-200V for 25-35 minutes. For quantification of NDUFA8 and MPC1, protein lysates were prepared in 1X Novex Tricine Sample Buffer (Thermo Fisher, LC1676) with 1X NuPAGE Sample Reducing Agent (Thermo Fisher, NP0009) and run on a Novex 10-20% Tricine Gel (Thermo Fisher, EC66255) in 1X Novex Tricine SDS Running Buffer (Thermo Fisher, LC1675) at constant 120V for 90 min. For quantification of PC, protein lysates were prepared in 1X Nu-PAGE LDS buffer (Thermo Fisher NP0007) with 1X NuPAGE reducing agent (Thermo Fisher NP0009) and run on a 3-8% Tris-Acetate gel (Thermo Fisher EA03755) in 1X Tris-acetate SDS running buffer (Thermo Fisher LA0041). PageRuler Plus/PageRuler Prestained Protein Ladder (Thermo Fisher, 26619 or 26616) was used as a molecular weight standard. Following PAGE, proteins were transferred onto nitrocellulose membranes (Thermo Fisher, IB23002) using an iBlot2 dry transfer system (Thermo Fisher) following manufacturer’s guidelines. Membranes were trimmed and blocked in 5% non-fat dry milk reconstituted in 1X TBS/0.1% Tween-20 (TBST) for 30min-2hr at room temperature with rocking. Blots were probed with primary antibodies diluted in 5% bovine serum albumin (Sigma, A4503) in 1X TBST overnight in a 4C cold room with rocking. The next day, blots were washed three times with 1X TBST, incubated in IRDye 680RD Goat anti-Rabbit and/or IRDye 800CW Goat anti-Mouse secondaries (Li-Cor) diluted in 1X TBST, washed three more times with 1X TBST, and imaged on a LiCor Odyssey Near-Infrared scanner (Li-Cor). Western blots were edited and exported using Image Studio software (Li-Cor) and densitometry was performed in Fiji. Refer to the Key Resources Table for specific information regarding antibody sources and dilutions.

### Intracellular metabolite quantification using liquid chromatography-mass spectrometry (LCMS)

For LCMS measurements, cells were seeded at a density of 75-200K cells/well of a 6-well plate. The following day, media was changed into the appropriate media (along with any indicated drug treatments/supplements) and cells were returned to the tissue culture incubator for the indicated times (6 or 24 hours). To extract intracellular polar metabolites, cells were washed 2-3 times with ice-cold blood bank saline (Fisher, 23293184), scraped in 500*µ*l of 80% HPLC-grade methanol (Fisher, A452SK) in HPLC-grade water (Sigma, 270733) and transferred to Eppendorf tubes. Samples were centrifuged (17,000g, 15min, 4C) to pellet insoluble material, and 400 *µ*L of each supernatant was transferred to a new centrifuge tube and lyophilized at 4C in a refrigerated vacuum centrifuge (CentriVap). Each treatment condition was plated in duplicate, and duplicate wells were trypsinized at the same time as metabolite extraction and counted on a Beckman Coulter Counter to obtain total cell volume per well. Average cell volumes for each condition were calculated, and lyophilized samples were resuspended at a constant resuspension volume: cell volume ratio (typically, 50-60*µ*l resuspension volume: 1*µ*l total cell volume) in 80% HPLC-grade methanol containing either of two 13C-labelled metabolite standards [U-^13^C-labeled yeast extract (Cambridge Isotope Laboratories, ISO1) or U-^13^C spirulina extract generated in-house by partial hydrolysis (12h in 6M HCl at 90C) of spirulina whole-cell lyophilized powder (Cambridge Isotope Laboratories, CLM-8400-PK)]. Most metabolite quantitation was performed using a Q Exactive HF-X Hybrid Quadrupole-Orbitrap Mass Spectrometer equipped with an Ion Max API source and H-ESI II probe, coupled to a Vanquish Flex Binary UHPLC system (Thermo Scientific). Mass calibrations were completed at a minimum of every 5 days in both the positive and negative polarity modes using LTQ Velos ESI Calibration Solution (Pierce). Polar Samples were chromatographically separated by injecting a sample volume of 1*µ*l into a SeQuant ZIC-pHILIC Polymeric column (2.1×150mm, 5mM, EMD Millipore). The flow rate was set to 150*µ*l/min, the autosampler temperature set to 10C and column temperature set to 30C. Mobile Phase A consisted of 20mM ammonium carbonate and 0.1% (v/v) ammonium hydroxide and Mobile Phase B consisted of 100% acetonitrile. The sample was gradient eluted (%B) from the column as follows: 0–20min: linear gradient from 85% to 20% B; 20–24min: hold at 20% B; 24–24.5min: linear gradient from 20% to 85% B; 24.5min–end: hold at 85% B until equilibrated with ten column volumes. Mobile phase was directed into the ion source with the following parameters: sheath gas = 45, auxiliary gas = 15, sweep gas = 2, spray voltage = 2.9kV in the negative mode or 3.5kV in the positive mode, capillary temperature = 300C, RF level = 40%, auxiliary gas heater temperature = 325C. Mass detection was conducted with a resolution of 240,000 in full-scan mode, with an AGC target of 3,000,000 and maximum injection time of 250ms. Metabolites were detected over a mass range of 70–1,050m/z. Quantitation of all metabolites was performed using Tracefinder v.4.1 (Thermo Scientific) referencing an in-house metabolite standards library using ≤ 5ppm mass error. For metabolite measurement in MPC-, PC-, and GOT1/2-KO cells, polar metabolite extracts were generated as described above and resuspended in HPLC-grade 80% methanol without stable isotope standards.

Metabolite quantitation was performed using an Agilent 6495D Triple-Quadrupole Mass Spectrometer equipped with an Agilent JetStream Heated ESI source coupled to an Agilent 1290 Infinity II UHPLC (Agilent Technologies). A Checktune was performed using the Agilent ESI-L low concentration tuning mix to assess the status of the instrument before data collection. Polar Samples were chromatographically separated by loading a sample volume of 2*µ*l onto one of two Poroshell 120 HILIC-Z columns (2.1×150mm, 2.7*µ*m, Agilent Technologies) running in a bespoke alternating column regeneration method. The flow rate was set to 600*µ*l/min, the multisampler temperature set to 4C and column temperature set to 25C. Mobile phase A consisted of 0.1% formic acid with 10mM ammonium formate and mobile phase B consisted of acetonitrile with formic acid at 0.1% (v/v). The sample was gradient eluted (%B) from the column as follows: 0–0.14min, initial hold at 95% ‘B’; 0.14–2.29min, 95% to 40% ‘B’; 2.29–4.0min and hold at 40% ‘B’. At the end of the method, the column oven valve would switch over to the other (K’ value matched) HILIC-Z column and the gradient would run on column 2, while column 1 was regenerated at 40% to 95% ‘B’ from 0–0.56min, followed by an increase in flow rate to 1,200*µ*l/min and held at 95% ‘B’ until ten column volumes were pumped through the column. Samples were analyzed on the mass spectrometer using the following parameters: gas flow = 13.0 l/min, sheath gas = 11 l/min, nebulizer = 35psi, gas temperature = 200C, sheath gas temperature = 250C, capillary voltage = 3,000V, nozzle voltage = 1,500V and a CAV voltage of 5V. Metabolites were targeted in the multiple reaction monitoring mode with a dwell time of 20ms at unit resolution; compound collision energies were previously optimized in the automated mode of MassHunter’s built in optimizer module (v.12.1). Quantitation of all metabolites was performed using MassHunter’s Quantitative Analysis module referencing an in-house metabolite standards library. Whenever possible, ion counts for metabolites of interest were normalized to the ^13^C-labeled version of that metabolite to account for matrix and loading effects. Statistical analysis was conducted using GraphPad Prism version 10.

### Stable isotope tracing

For stable isotope tracing experiments, cells were plated as above. The following day, cells were washed twice with DPBS and changed to media with the appropriate treatments and tracer. For glutamine tracing, cells were switched into DMEM without glucose, glutamine, pyruvate or phenol red (Sigma, D5030) supplemented with 10% FBS, 1% penicillin–streptomycin, 1mM pyruvate, 25mM 12C glucose (Sigma, G7528) and 4mM U-13C glutamine (Cambridge Isotopes, CLM-1822). For glucose tracing, 143B cells were switched into DMEM without glucose, glutamine, pyruvate or phenol red supplemented with 10% FBS, 1% penicillin–streptomycin, 1mM alpha-ketobutyrate (Sigma), 25mM U-^13^C glucose (Cambridge Isotopes, CLM-1396-PK) and 4mM U-12C glutamine (Sigma, G5792). 24 hours after tracer administration, polar metabolites were extracted and analyzed on a Q Exactive HF-X Hybrid Quadrupole-Orbitrap Mass Spectrometer as above, and data was analyzed as above using TraceFinder v4.1. Natural abundance corrections were performed using IcoCor v.2.2, incorporating the known isotopic purity of the specific tracers used.

### Cell cycle analysis

Cell cycle analysis was performed similar to what was previously described ^24^. Cells were seeded at 50-200K/well in a 12-well plate. The following morning, media was changed and cells were returned to the incubator. 48 hours later, cells were trypsinized, pelleted, washed twice, then resuspended in 300 *µ*L ice cold DPBS. While vortexing, 700 *µ*L ice-cold 100% ethanol (Decon, 2716) was added dropwise to fix cells. Fixed cells were stored at −20C until being processed for flow cytometry, which entailed pelleting the cells, washing twice with DPBS, then staining in 250*µ*L of a 50*µ*g/ml propidium iodide (Biotium, 40017) with 100*µ*g/ml RNase A (QIAGEN) staining solution for 1h. overnight at 4C, protected from light. Samples were then passed through a 0.35*µ*m filter into flow cytometry tubes (Falcon) before being analyzed on a BD FACSymphony A52 Cell Analyzer running FACSDiva software. 10,000 events were recorded per sample. Data were analyzed using the ‘Cell Cycle’ analysis module of FlowJo v.10.10.1.

### DepMap analysis

All data for DepMap analyses were pulled from the publicly available 26Q1 dataset using the Data Explorer (https://depmap.org/portal/data_explorer_2/). For each queried gene, the data was filtered to remove cell lines with missing values for either the queried gene or average SDH dependency (calculated using the ‘mean’ of a multi-gene query functionality in Data Explorer). For each gene, cell lines were filtered to those with the top and bottom 5% of expression, and mean SDH dependency scores were calculated and compared between the two groups. Data were manipulated using R v4.4.0 and statistics (including regressions) were calculated/plotted using GraphPad Prism version 10.

### Statistics and reproducibility

Unless otherwise noted, data was manipulated in Microsoft Excel version 16.108.3, and all graphs and statistical analyses were made in GraphPad Prism version 10. Replicates, defined as parallel biological samples independently treated, collected and analyzed during the same experiment, are shown. Experiments were verified with two or more independent repetitions showing qualitatively similar results. Details pertaining to all statistical tests can be found in the figure legends. No statistical test was used to predetermine sample sizes, but our sample sizes are similar to those reported in previous publications ^15,24^. The investigators were not blinded to treatment allocation during experiments and outcome assessment. The experiments were not randomized. Data distribution was assumed to be normal, but this was not formally tested.

## ACKNOWLEDGMENTS

We thank members of the Sullivan laboratory and J.C. Young for discussion and feedback. This research was supported by the Proteomics & Metabolomics and Flow Cytometry Shared Resources of the Fred Hutchinson Cancer Center/University of Washington Cancer Consortium (P30CA015704). L.B.S. acknowledges support from the National Institute of General Medical Sciences (NIGMS) (R35GM147118) and the National Cancer Institute (U54CA132381). D.S. was supported by a Public Health Services Ruth L. Kirschstein National Research Service Award (T32GM007270) from NIGMS and a Paul Neiman Graduate Student Award. The funders had no role in study design, data collection and analysis, decision to publish or preparation of the manuscript. This LATEXdocument was created in Overleaf and adapted from the Interdisciplinary Laboratory of Computational Social Science (iLCSS) template.

## Notes

### Competing Interest Statement

The authors have declared no competing interest.

### Summary of Updates

During preparation of the source data, it was discovered that the UMP abundance data in Figure S6I was incorrect. This figure panel has been revised, and the overall conclusions of the figure/work are unaffected.

