## Supplement for "Adaptive plasticity of aspartate metabolism in succinate dehydrogenase-deficient cancer cells"

### Supplemental information for: adaptive plasticity of aspartate metabolism in succinate dehydrogenase-deficient cancer cells

**David Sokolov**[1], **Eric Zheng**[1], **Serwah Danquah**[1], **Madeleine L. Hart**[1], **and Lucas B. Sullivan**[1,2]

[1] Human Biology Division, Fred Hutchinson Cancer Center, Seattle, WA, 98109, USA.

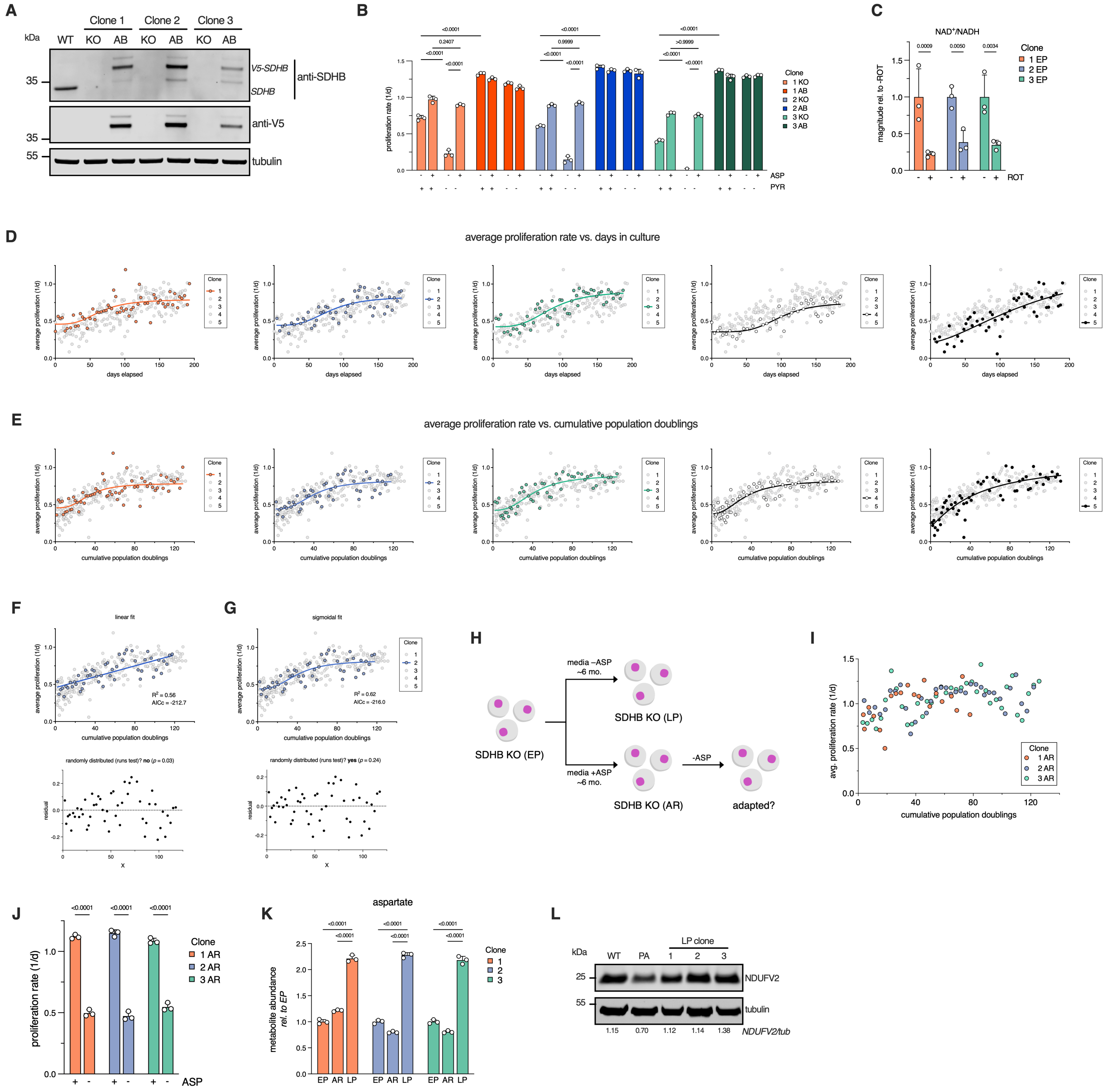


**Fig. S1. Related to Figure 1.** (**A**) Representative western blot showing levels of SDHB, V5-tagged SDHB, and tubulin loading control in wild-type parental cells (WT), early passage (EP), and addback (AB) SDHB knockout clones 1-3. (**B**) Absolute proliferation rates (mean +/- S.D.) of early passage (EP) and addback (AB) clones 1-3 in DMEM with all combinations of 1 mM pyruvate and 20 mM aspartate supplementation. Statistical significance determined using an ordinary two-way ANOVA and Tukey's multiple comparisons test with a single pooled variance. Data for KO clones in pyruvate-containing media are the same as plotted in Figure 1H. (n=3) (**C**) Relative whole-cell $NAD^+$/NADH (mean +/- S.D.), measured using LCMS, in SDHB-KO clones 1-3 after 6 hours of treatment with vehicle control or 50 nM rotenone (ROT). Values are normalized to the vehicle-treated condition in each respective clone. Statistical significance determined using an ordinary two-way ANOVA and uncorrected Fisher's LSD with a single pooled variance. (n=3) (**D-E**) Average inter-passage proliferation rates (see methods) of individual knockout clones plotted against elapsed calendar days (**D**) or cumulative population doublings (**E**) fit with sigmoidal curves. (**F-G**) Linear (**F**) and sigmoidal (**G**) fits of knockout clone 2 average proliferation rate data. $R^2$ and Akaike information criterion (AICc) is shown, and residuals are plotted beneath the fit. Results of runs test for residual clustering are shown above the residual plot. (**H**) Schematic illustrating the creation of aspartate reared (AR) variants of knockout clones, which are cultured long-term in media supplemented with 20 mM aspartate, before aspartate is withdrawn and cells are evaluated for potential adaptation. (**I**) Average inter-passage proliferation rates (see methods) of aspartate reared (AR) clones 1-3 over 80-120 cumulative population doublings. (**J**) Absolute proliferation rates (mean +/- S.D.) of aspartate-reared (AR) SDHB knockout clones 1-3 with or without 20 mM aspartate supplementation. Statistical significance determined using an ordinary two-way ANOVA and uncorrected Fisher's LSD with a single pooled variance. (n=3) (**K**) Relative whole-cell aspartate levels (mean +/- S.D.) of early passage (EP), aspartate reared (AR), and late passage (LP) SDHB knockout clones 1-3. Levels are normalized to each corresponding EP clone. Statistical significance determined using an ordinary two-way ANOVA and Tukey's multiple comparisons test with a single pooled variance. (n=3) (**L**)Representative western blot showing levels of NDUFV2 and tubulin loading control in wild-type parental cells (WT), a previously adapted (PA) SDHB-KO clone that adapted by suppressing complex I [?] , and late passage (LP) SDHB knockout clones 1-3. Normalized NDUFV2 band densities are shown below each respective lane. Statistical significance determined using ordinary two-way ANOVAs and uncorrected Fisher's LSD with single pooled variance (**C,J**) or ordinary two-way ANOVA and Tukey's multiple comparisons test with a single pooled variance (**B,K**).

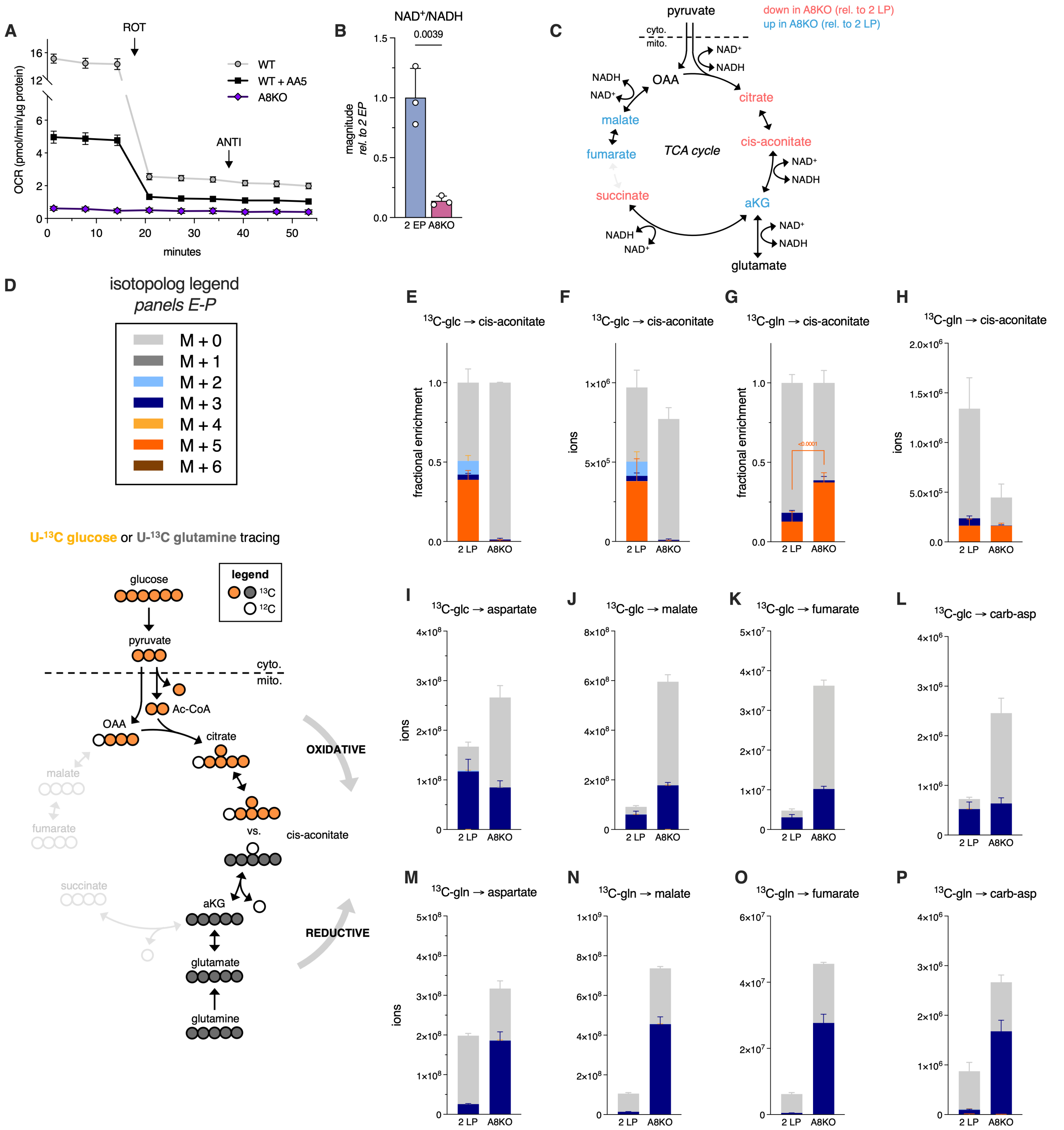

**Fig. S2. Comparing adapted metabolic states of CI-high and CI-low SDH-deficient cells, related to Figure 2.** (**A**) Normalized oxygen consumption rate (OCR) traces (mean +/- S.D.) for wild-type parental cells (WT), parental cells treated with 5 $\mu$M Atpenin A5 (WT + AA5), and early passage SDHB-KO NDUFA8-KO clone 2 (A8KO). Injections of rotenone (ROT) and antimycin (ANTI) are shown with arrows. (n=7-10) (**B**) Relative whole-cell $NAD^+$/NADH (mean +/- S.D.) in early passage SDHB KO clone 2 (2 EP) and A8KO 24 hours after media change. Levels are normalized to 2 EP. (n=3) (**C**) Schematic depicting TCA cycle metabolites which are more or less abundant in A8KO compared to late passage SDHB KO clone 2 (2 LP) (original data in Fig. 2E), highlighting redox-dependent interconversions. (**D**) Isotopolog legend for panels E-P and schematic illustrating U-$^{13}$C glucose/U-$^{13}$C glutamine tracing strategy to quantify oxidative vs. reductive TCA cycle metabolism in SDHB-KO cells. (**E-H**) Fractional (**E,G**) or absolute (**F,H**) isotopolog distributions (mean +/- S.D.) for cis-aconitate in 2 LP and A8KO following 24 hours of tracing in media containing U-$^{13}$C glucose (**E,F**) or U-$^{13}$C glutamine (**G,H**). In (**G**), p-value depicts the results of significance testing on the M+5 fraction. (n=3) (**I-P**) Absolute isotopolog distributions for the indicated metabolites (mean +/- S.D.) in late passage 2 LP and A8KO following 24 hours of tracing in media containing U-$^{13}$C glucose (**I-L**) or U-$^{13}$C glutamine (**M-P**). (n=3) Statistical significance determined using an unpaired t-test (**B**) or ordinary two-way ANOVA and Sidak's multiple comparisons test with a single pooled variance (**G**).

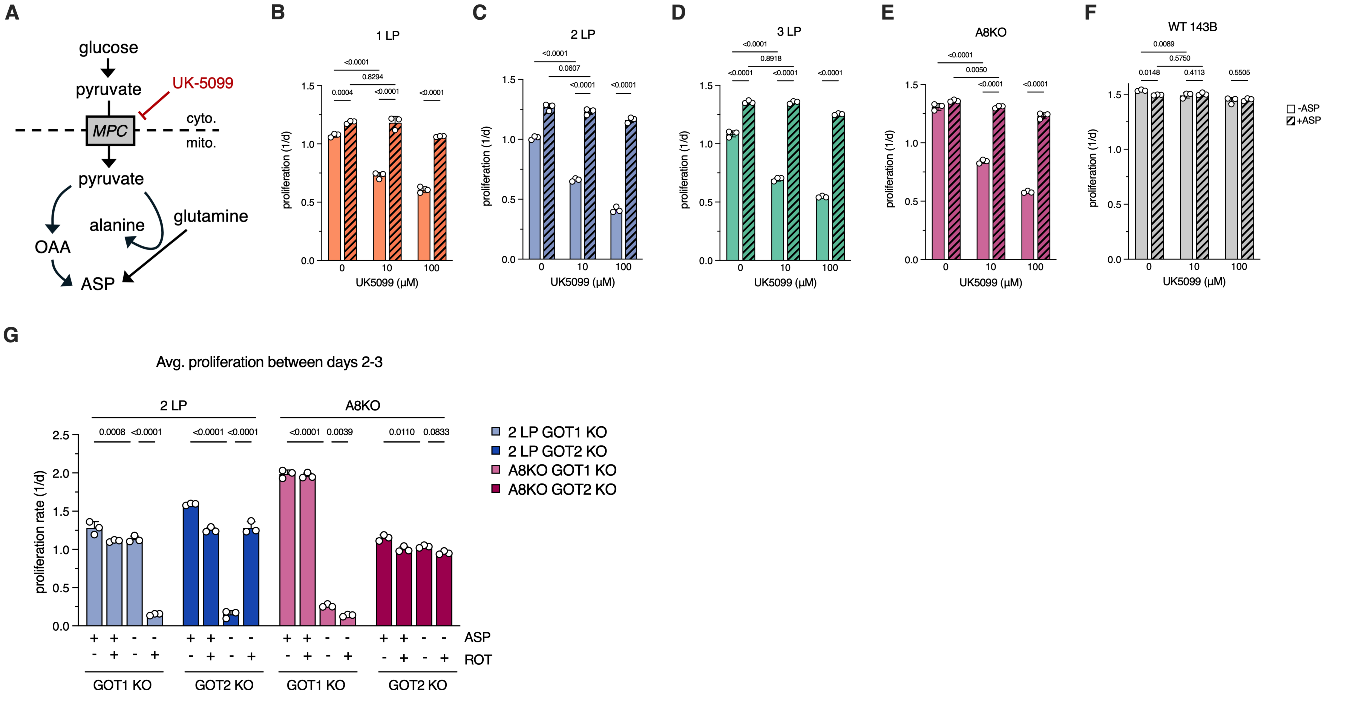


**Fig. S3. Related to Figure 3.** (**A**) Schematic illustrating the role of the mitochondrial pyruvate carrier (MPC) in alternative aspartate synthesis and its inhibition using UK-5099 (**B-F**) Proliferation rates (mean +/- S.D.) of late passage SDHB-KO clones 1-3 (1-3 LP) (**B-D**), A8KO (**E**), or wild-type 143B parental cells (143B) (**F**) following treatment with the indicated doses of UK-5099, with or without 20 mM aspartate supplementation. Statistical significance determined using an ordinary two-way ANOVA and uncorrected Fisher's LSD with a single pooled variance. (n=3) (**G**) Data from the same experiment as in Figure 3M, with proliferation rates (mean +/- S.D.) calculated between days 2-3 of the 3-day assay. Statistical significance determined using an ordinary two-way ANOVA and uncorrected Fisher's LSD with single pooled variance.

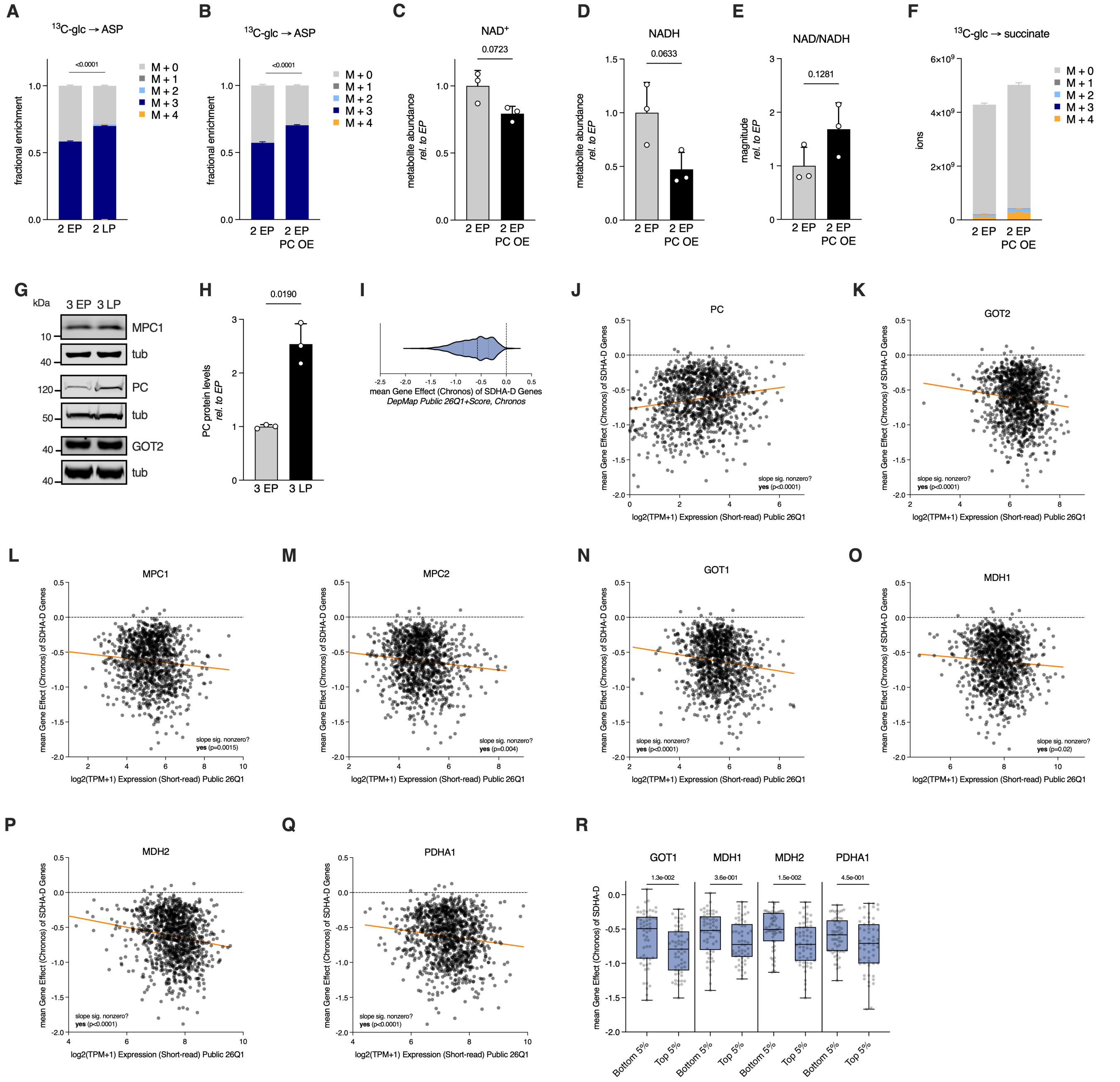


**Fig. S4. Related to Figure 4.** (**A**) Fractional isotopolog distributions (mean +/- S.D.) for aspartate in early/late passage (EP/LP) SDHB-KO clone 2 following 24 hours of tracing in media containing U-$^{13}$C glucose. (n=3) (**B**) Fractional isotopolog distributions (mean +/- S.D.) for aspartate in early passage SDHB-KO clone 2 (2 EP) and PC-overexpressing 2 EP following 24 hours of tracing in media containing U-$^{13}$C glucose. (n=3) (**C-E**) Relative NAD$^+$ (**C**), NADH (**D**), or NAD$^+$/NADH (**E**) levels (mean +/- S.D.) of 2 EP and 2 EP PC OE 24 hours after media change. Levels are normalized to 2 EP. (n=3) (**F**) Absolute isotopolog distributions (mean +/- S.D.) for succinate in 2 EP and 2 EP PC OE following 24 hours of tracing in media containing U-$^{13}$C glucose. (n=3) (**G**) Representative western blots showing levels of MPC1, PC, GOT2, and tubulin loading control in early passage (EP) and late passage (LP) SDHB KO clone 3. (**H**) Quantification of PC protein levels from 3 EP/LP using western blotting. (n=3) (**I**) Mean SDHA-D gene effect scores for all cell lines in DepMap 26Q1 (**J-Q**) Mean SDHA-D gene effect scores for cell lines in DepMap 26Q1 plotted against relative expression of PC (**J**), GOT2 (**K**), MPC1 (**L**), MPC2 (**M**), GOT1 (**N**), MDH1 (**O**), MDH2 (**P**), and PDHA1 (**Q**) fitted with linear regressions (n=1140 cell lines) (**R**) Mean SDHA-D DepMap gene effect scores for the top and bottom 5% of expressors of the indicated metabolic genes. (n=57) Statistical significance determined using Welch's t-test (**C-E, H**), or ordinary one-way (**R**) or two-way (**A-B**)ANOVAs with Sidak's multiple comparisons test with a single pooled variance. Whether the slope of the regression lines were significantly nonzero was determined using Prism.

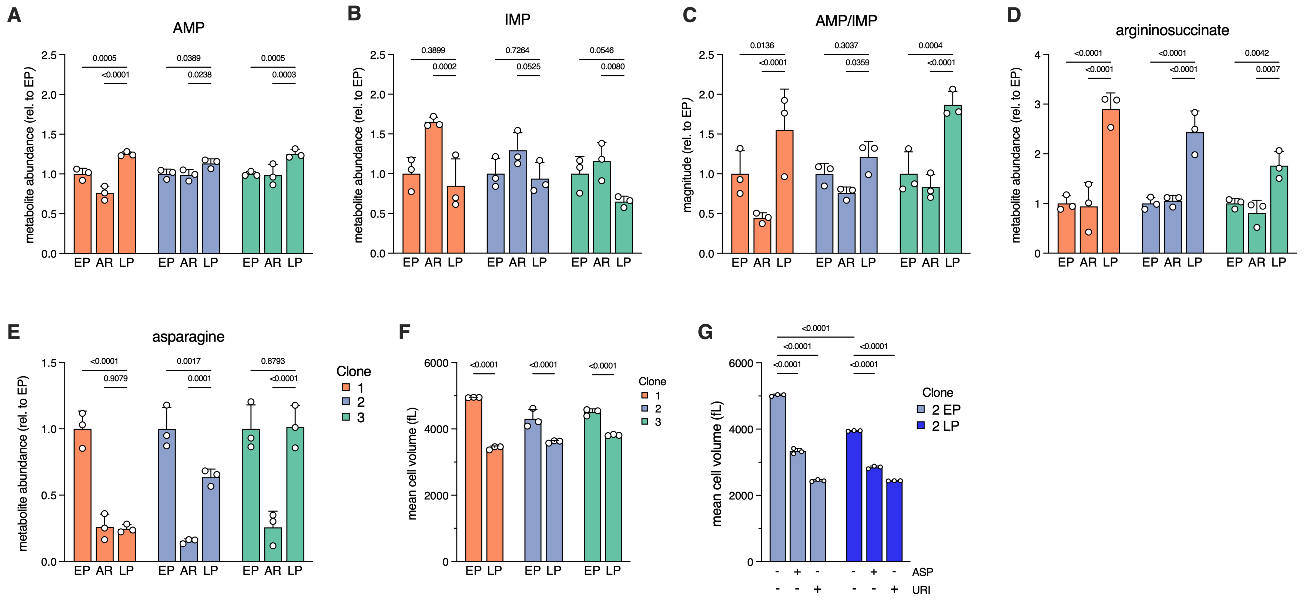


**Fig. S5. Related to Figure 5.** (**A-E**) Relative levels of adenosine monophosphate (AMP) (**A**), inosine monophosphate (IMP) (**B**), AMP/IMP (**C**), argininosuccinate (**D**), or asparagine (**E**) (mean +/- S.D.) in early passage (EP), aspartate reared (AR), and late passage (LP) SDHB knockout clones 1-3. Levels are normalized to each corresponding EP clone. (n=3) (**F**) Mean per-cell volumes of EP/LP SDHB KO clones 1-3. (n=3) (**G**) Mean per-cell volume of EP/LP SDHB KO clone 2 supplemented with 20 mM aspartate (ASP) or 500 $\mu$M uridine (URI). (n=3) Statistical significance determined using ordinary two-way ANOVAs and uncorrected Fisher's LSD with single pooled variance (**A-F**) or ordinary two-way ANOVAs and Tukey's multiple comparisons test with single pooled variance (**G**).

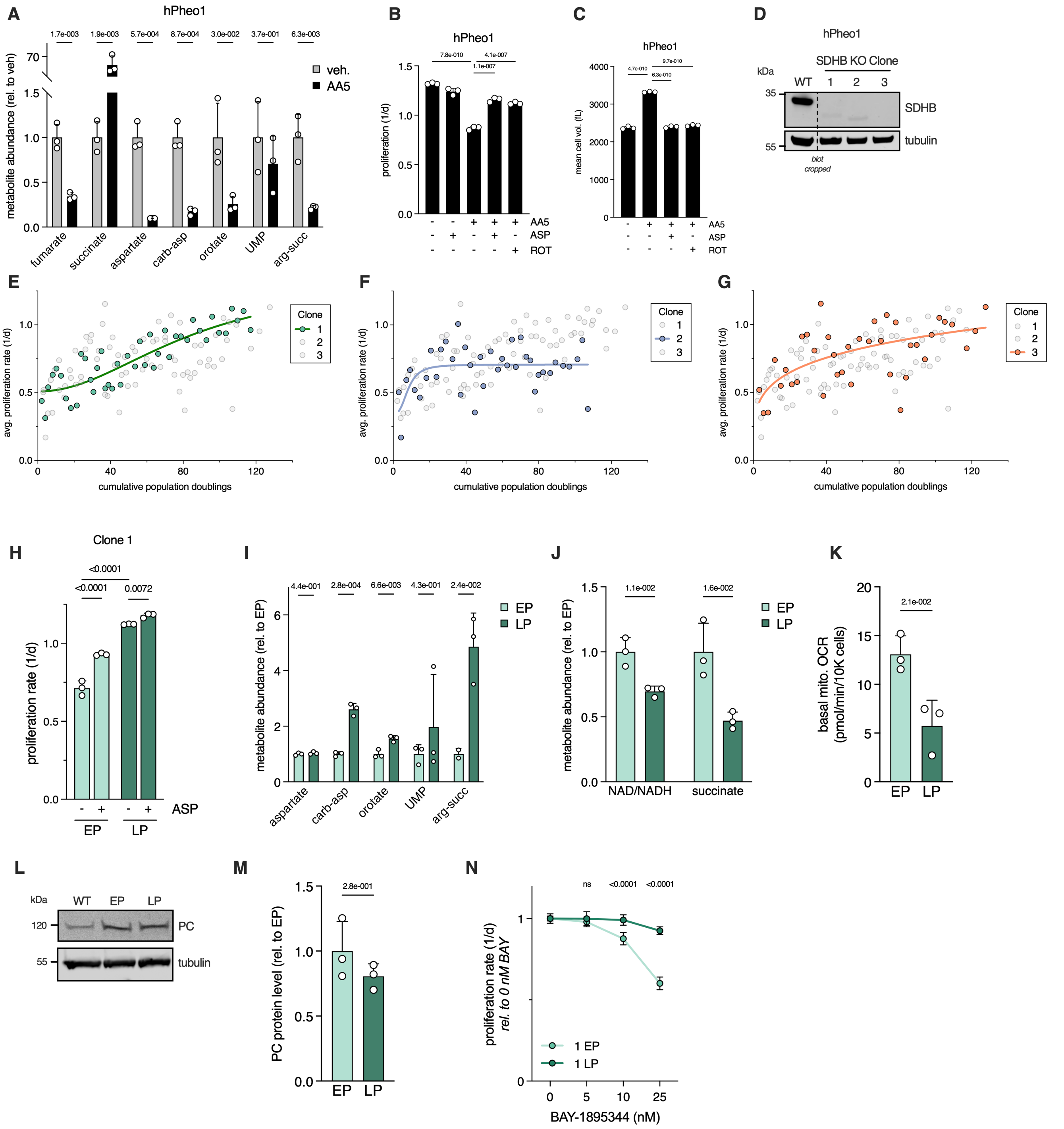


**Fig. S6. Adaptation in SDH-deficient human pheochromocytoma cells.** (**A**) Relative abundance (mean +/- S.D.) of the indicated metabolites in hPheo1 cells treated with 5 $\mu$M AA5 or vehicle control for 24 hours. Metabolite abundances are normalized to vehicle-treated controls. (n=3). (**B**) Proliferation rates (mean +/- S.D.) of hPheo1 cells treated with the indicated combinations of 5 $\mu$M AA5, 20 mM aspartate (ASP), and 100 nM rotenone (ROT). (n=3). (**C**) Mean per-cell volume (mean +/- S.D.) of hPheo1 cells with the indicated combinations of 5 $\mu$M AA5, 20 mM aspartate (ASP), and 100 nM rotenone (ROT). (n=3). (**D**) Western blot showing expression of SDHB and tubulin loading control in wild-type (WT) hPheo1 cells and three hPheo1 SDHB knockout clones. (**E-G**) Average inter-passage proliferation rates (see methods) of hPheo1 SDHB KO clones 1-3 over 120 cumulative population doublings, with overlaid sigmoidal curve fit. (**H**) Proliferation rates (mean +/- S.D.) of early passage (EP) and late passage (LP) hPheo1 SDHB KO clone 1 with or without 20 mM aspartate (ASP) supplementation. (n=3). (**I**) Relative abundance (mean +/- S.D.) of the indicated metabolites in EP/LP hPheo1 SDHB KO clone 1. Metabolite abundances are normalized to EP. (n=3). (**J**) Relative magnitude (mean +/- S.D.) of $NAD^+$/NADH and succinate abundance in EP/LP hPheo1 SDHB KO clone 1. Metabolite abundances are normalized to EP. (n=3). (**K**) Basal mitochondrial oxygen consumption rates (mito. OCR) (mean +/- S.D.) of EP/LP hPheo1 SDHB KO clone 1. (n=3). (**L**) Western blot showing expression of PC and tubulin loading control in wild-type (WT) hPheo1 cells and EP/LP hPheo1 SDHB KO clone 1 (**M**) PC protein levels (mean +/- S.D.) in EP/LP hPheo1 SDHB KO clone 1 determined using western blotting of replicate samples and densitometry. (n=3). (**N**) Proliferation rates (mean +/- S.D.) of EP/LP hPheo1 SDHB knockout clone 1 in the indicated doses of BAY-1895344. Proliferation rates are normalized to the 0 nM BAY dose in each condition. P-values shown correspond to statistical testing comparing the relative proliferation rate of EP vs. LP at the indicated dose of BAY. (n=3) Statistical significance determined using multiple unpaired t-tests (**A,I-J**), ordinary one-way (**B-C**) or two-way (**H**) ANOVAs and Tukey's multiple comparisons test with single pooled variance (**B-C**), ordinary two-way ANOVAs and an uncorrected Fisher's LSD with a single pooled variance (**N**), or Welch's t-tests (**K,M**).

| Clone | Bottom (doublings/day) | Top (doublings/day) | Span (doublings/day) | IC50 (doublings) | $R^2$ |
|---|---|---|---|---|---|
| 1 | 0.45 | 0.78 | 0.32 | 33.75 | 0.50 |
| 2 | 0.44 | 0.82 | 0.38 | 39.87 | 0.62 |
| 3 | 0.42 | 0.89 | 0.47 | 41.36 | 0.74 |
| 4 | 0.37 | 0.83 | 0.45 | 33.57 | 0.77 |
| 5 | 0.20 | 1.09 | 0.88 | 46.54 | 0.73 |
| avg. | 0.37 | 0.882 | 0.5 | 39.01 | |

**Supplemental Table 1. Table of fitted curve parameters for each knockout clone (plotted in Figure S1E) and the average of all five clones, including bottom (doublings/day), top (doublings/day), span (top-bottom, doublings/day), 'IC50' (the number of cumulative population doublings needed to achieve half of the 'top' proliferation rate), and $R^2$.**

**Key Resources Table**

| Resource/Reagent | Manufacturer | Catalog Number | Sequence (5'-3') if applicable | Dose/dilution used |
|---|---|---|---|---|
| SDHB sgRNA 1 | Synthego | N/A | UCGCCCUCUCCUUGAGGCG | See Methods |
| SDHB sgRNA 2 | Synthego | N/A | AGAAAUUUGCCAUCUAUCGA | See Methods |
| NDUFA8 sgRNA 1 | Synthego | N/A | CACAUUGAGCUCCAUAGUGA | See Methods |
| NDUFA8 sgRNA 2 | Synthego | N/A | AAAGCUGCGGCCCAUCACUA | See Methods |
| NDUFA8 sgRNA 3 | Synthego | N/A | UUCUGCUGUGCUUAAAGCUG | See Methods |
| MPC1 sgRNA 1 | Synthego | N/A | UUGCCUACAAGGUACAGCCU | See Methods |
| MPC1 sgRNA 2 | Synthego | N/A | GGGCUACUUCAUUUGUUGCG | See Methods |
| MPC1 sgRNA 3 | Synthego | N/A | AUGUCAAAGAAUAGCAACAG | See Methods |
| PC sgRNA 1 | Synthego | N/A | GCAGGCCCGGAACACACGGA | See Methods |
| PC sgRNA 2 | Synthego | N/A | GCUGGAGGAGAAUUACACCC | See Methods |
| PC sgRNA 3 | Synthego | N/A | ACACCGGCCGCAUUGAGGU | See Methods |
| GOT1 sgRNA 1 | Synthego | N/A | CAGUCAUCCGUGCGAUAUGC | See Methods |
| GOT1 sgRNA 2 | Synthego | N/A | GCACGGAUGACUGCCAUCCC | See Methods |
| GOT1 sgRNA 3 | Synthego | N/A | CGAUCUUCUCCAUCUGGGAA | See Methods |
| GOT2 sgRNA 1 | Synthego | N/A | UUUCUCAUUUCAGCUCCUGG | See Methods |
| GOT2 sgRNA 2 | Synthego | N/A | CGGACGCUAGGCAGAACGUA | See Methods |
| GOT2 sgRNA 3 | Synthego | N/A | UCCUUCCACUGUUCCGGACG | See Methods |
| Aspartate | Sigma | A7219 | N/A | 20 mM |
| Uridine | Sigma | U3003 | N/A | 500 $\mu$M |
| UK-5099 | Cayman Chemical | 16980 | N/A | Varies, see figure |
| Rotenone | Sigma | R8875 | N/A | 50-100 nM |
| Antimycin | Sigma | A8674 | N/A | 500 nM |
| Anti-SDHB | Atlas Antibodies | HPA002868 | N/A | 1:1000 |
| Anti-tubulin | Sigma | T6199 | N/A | 1:3000 |
| Anti-V5 | Cell Signaling | 13202S | N/A | 1:2000 |
| Anti-NDUFV2 | Proteintech | 15301-1-AP | N/A | 1:1000 |
| Anti-NDUFA8 | Atlas Antibodies | HPA041510 | N/A | 1:1000 |
| Anti-MPC1 | Cell Signaling | 14462S | N/A | 1:1000 |
| Anti-PC | Proteintech | 66615-1 | N/A | 1:1000 |
| Anti-GOT2 | Proteintech | 14800-1-AP | N/A | 1:1000 |
| Anti-GOT1 | Cell Signaling | 34423S | N/A | 1:1000 |
| pLX304_PC | DNASU Plasmid Repository | HsCD00436386 | N/A | N/A |
| pLenti6.3-V5-DEST_SDHB | DNASU Plasmid Repository | HsCD00954622 | N/A | N/A |
| pLenti6.3-V5-DEST_MPC1 | DNASU Plasmid Repository | HsCD00942383 | N/A | N/A |
| BAY-1895344 | Selleckchem | S8666 | N/A | Varies, see figure |
